# Evoked Resonant Neural Activity Long-Term Dynamics can be Reproduced by a Computational Model with Vesicle Depletion

**DOI:** 10.1101/2024.02.25.582012

**Authors:** James J. Sermon, Christoph Wiest, Huiling Tan, Timothy Denison, Benoit Duchet

**Affiliations:** Institute of Biomedical Engineering, Department of Engineering Science, University of Oxford, Oxford, UK; MRC Brain Networks Dynamics Unit, Nuffield Department of Clinical Neurosciences, University of Oxford, Oxford, UK

## Abstract

Subthalamic deep brain stimulation (DBS) robustly generates high-frequency oscillations known as evoked resonant neural activity (ERNA). Recently the importance of ERNA has been demonstrated through its ability to predict the optimal DBS contact in the subthalamic nucleus in patients with Parkinson’s disease. However, the underlying mechanisms of ERNA are not well understood, and previous modelling efforts have not managed to reproduce the wealth of published data describing the dynamics of ERNA. Here, we therefore aim to present a minimal model capable of reproducing the characteristics of the slow ERNA dynamics published to date. We make biophysically-motivated modifications to the Kuramoto model and fit its parameters to the slow dynamics of ERNA obtained from data. We further validate the model against experimental data from Parkinson’s disease patients by simulating variable stimulation and medication states, as well as the response of individual neurons. Our results demonstrate that it is possible to reproduce the slow dynamics of ERNA with a single neuronal population, and, crucially, with vesicle depletion as the key mechanism behind the ERNA frequency decay. We provide a series of predictions from the model that could be the subject of future studies for further validation.

**Author Summary:** ERNA is a high amplitude response to stimulation of deep brain structures, with a frequency over twice that of the frequency of stimulation. While the underlying mechanisms of ERNA are still unclear, recent findings have demonstrated its importance as the best indicator of which stimulation contact to select for DBS therapy in patients with Parkinson’s disease. Previous modelling studies of ERNA focus on the immediate responses to stimulation (*<*200ms) and rely on interconnected neural structures and delays. Our work shows that the long-term (on the scale of one or more seconds) ERNA response to continuous stimulation can be modelled using a single neural structure. The proposed model also captures the long-term frequency and amplitude characteristics of ERNA with variable stimulation and medication paradigms. The key features of the model, in particular the depletion of vesicles carrying neurotransmitters between neurons by high-frequency stimulation, may provide insights into the underlying mechanisms of ERNA and inform future investigations into this neural response.

## Introduction

Deep brain stimulation (DBS) of the subthalamic nucleus (STN) has been a widely used treatment for Parkinson’s disease (PD) for over two decades and has proven effective for the alleviation of Parkinsonian motor symptoms. During STN-DBS, evoked resonant neural activity (ERNA) is commonly observed in simultaneous recordings of the STN. However, the underlying mechanisms of this phenomenon are not well understood.

ERNA is a series of high-frequency oscillations (HFOs, *>*200Hz) of high-amplitude in response to 70-180Hz stimulation, despite there being very little to no high frequency activity at the stimulation site pre-stimulation [1] (also see 4 in Fig 2A). ERNA has been observed in different neural structures, but so far only in the pallidal [2, 3] and subthalamic regions [1, 4–6]. Furthermore, ERNA has also been characterised as a robust response across different devices, patient groups and stimulation conditions [1, 4–6]. However, the reproducibility and unphysiological appearance of ERNA have led to questions of whether it is of biophysical origin. The biophysical origin of ERNA was later demonstrated in [5], but many questions regarding the mechanisms of ERNA still remain.

Since Sinclair et al. observed the presence of ERNA in the Parkinsonian STN [4], several hypotheses and models looking to explain the presence of ERNA have been proposed. These hypotheses range from inhibition and/or excitation of neural ensembles to recruitment of inter-component projections between the STN and GPe [6–9]. However, thus far none of these propositions have managed to explain all ERNA characteristics, in particular the long term variation of ERNA with continuous DBS. Recently, the significance of comprehending the fundamental mechanisms of ERNA has become more apparent with evidence that ERNA may provide the strongest predictor of clinical performance for STN-DBS contact selection [10].

In this study, we propose a biophysically-motivated computational model that is capable of reproducing the key long-term ERNA characteristics currently described in the literature under a variety of stimulation paradigms, with a view to hypothesise the fundamental mechanisms responsible for ERNA. We assembled a single population of Kuramoto oscillators intended to represent the STN. Through the addition of synaptic vesicle depletion and a second-order coupling function to the standard Kuramoto model, we were able to fit the parameters of the modified Kuramoto model to key features of ERNA. We simulate further stimulation paradigms to validate and gain insights into the key properties of the computational model. In addition, we suggest predictions that can be used to further validate the computational model.

## Results

We modified the Kuramoto model to include a second-order coupling function as well as the effect of vesicle depletion from high-frequency DBS (see Methods sections for details), and fitted the modified model to salient long-term features of ERNA observed in patient data with DBS. By comparing simulations of the fitted model to ERNA frequency responses recorded in patients under varying stimulation frequencies, and with minimal additional parametrisation under varying stimulation amplitudes as well as medication states, we characterise the model and test its predictive power. We provide validation for the model by comparing the model’s ERNA amplitude response (which is not considered in the parameter tuning) to that of the data. Lastly, we perform simulations of the model analogous to in vitro and in vivo STN recordings to further test the model against biophysical data.

### Results of Fitting Process

The best-ranked parameter set resulting from the fitting to the peak frequency ERNA decay while limiting off-stimulation high-frequency activity (see Methods section *Fitting Process*) provided a good approximation of the peak frequency ERNA decay estimated from data recorded in a PD patient with 130Hz, 3mA STN DBS (Fig 1C). The corresponding model parameters are given in Table A in Supplementary Material. In the spectrogram, the fitted model shows a broad ERNA decay approximating patient data (compare Fig 1A and Fig 1B). Before stimulation onset, high-amplitude and high-frequency activity is absent in the model, with a lower amplitude peak at around 40Hz as in the data (compare Fig 1A and characteristic 3 in Fig 2A).

**Fig 1.**
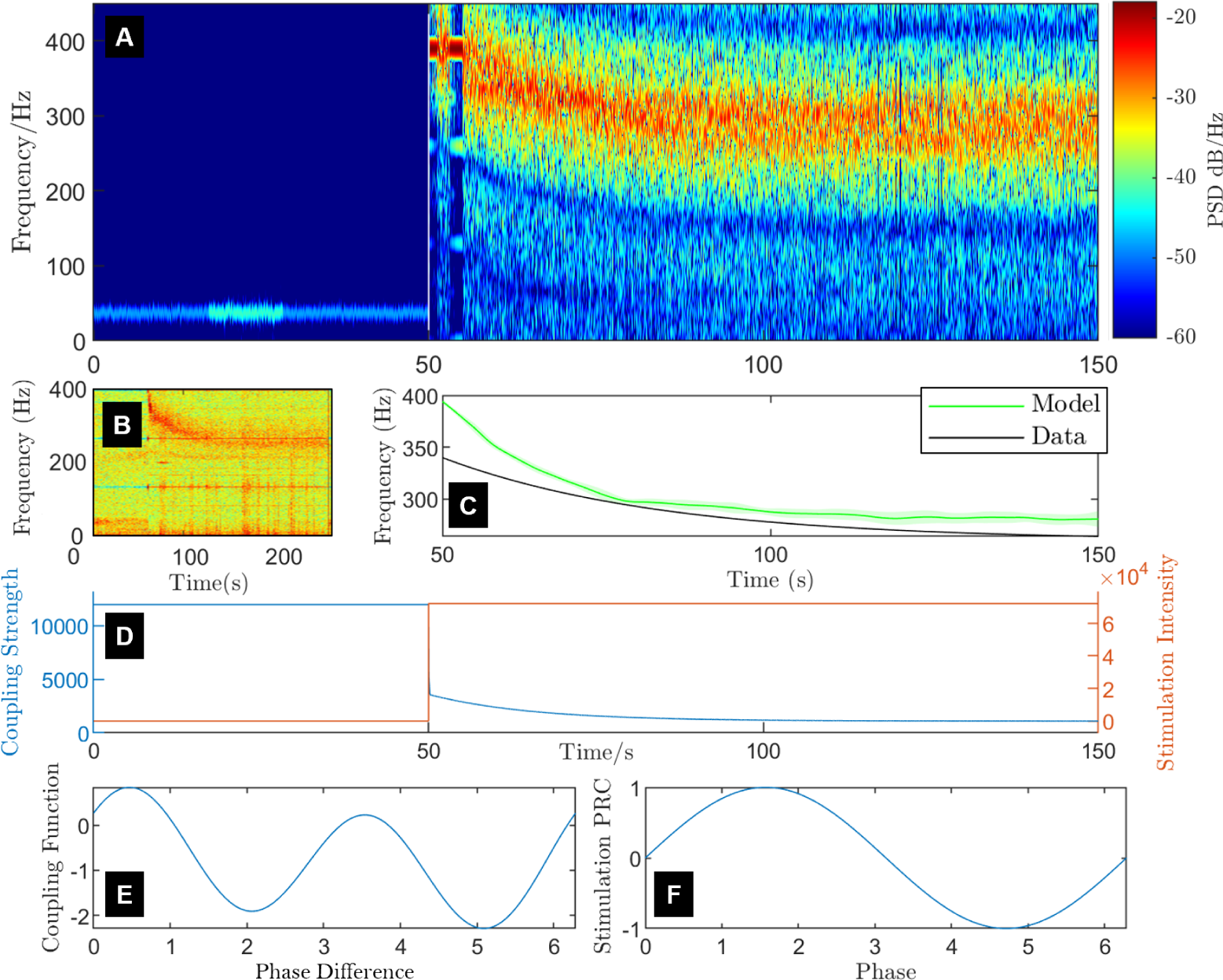
Fitted slow dynamics of the modified Kuramoto model on and off stimulation and model’s key properties. (A) Spectrogram showing the fitted model’s ERNA response to 130Hz stimulation (fitted stimulation amplitude). The transition from off to on stimulation happens at 50 seconds. The colour scale represents the power spectral density (PSD). (B) Example of the slow ERNA dynamics in the frequency domain for 130Hz and 3mA stimulation adapted from [1]. (C) Comparison of the peak ERNA frequency decay estimated from panel B and fed into the fitting process (black), and the model predicted ERNA frequency peak (green) averaged over 15 realisations of noise, natural frequencies and initial conditions for the 100 second period of stimulation following 50 seconds of the model settling off-stimulation. We also display the standard error of the mean for the averaged model output in the lighter green band. (D) Evolution of coupling strength in the model (blue) and stimulation intensity (orange) over the full stimulation window. Stimulation is a pulse train at 130Hz, but intensity is depicted as constant following stimulation onset in this figure for demonstrative purposes. (E) The fitted second-order Fourier series coupling function, and (F) the phase-response curve (PRC, here a sine wave) used in the model.

**Fig 2.**
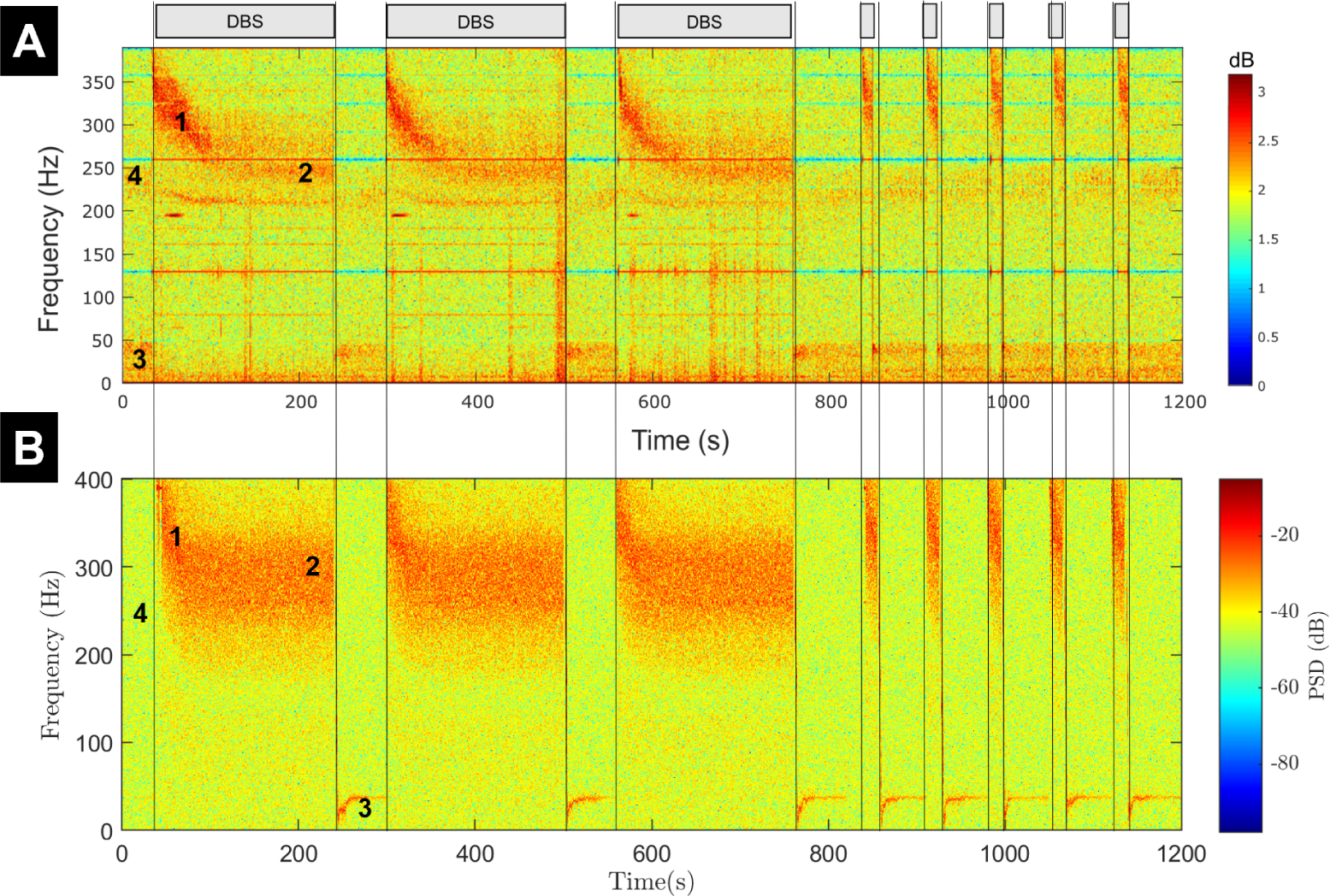
The modified Kuramoto model replicates ERNA characteristics observed in data during multiple on-DBS windows with pauses. (A) Data spectrogram, adapted from [1]. On-DBS periods (with 130Hz, 3mA STN DBS) are indicated by grey rectangles. (B) Spectrogram from the modified Kuramoto model with added white noise. Both panels use similar time points of DBS initiation and termination. Annotations are described in the main text. Since the model output is on a very different scale from the data, the scale of the model output PSD differs from that of the data.

The first key modification of the Kuramoto model that was found to be necessary for reproducing the slow ERNA dynamics is the reduction of coupling strength through vesicle depletion during stimulation (Fig 1D). Over the timescale during which the slow ERNA dynamics develop, the readily-releasable vesicle pool (RRP) as well as the recycling pool (RP) are very quickly exhausted, and the resting pool (RtP) becomes the dominant vesicle pool for synaptic transmission [11–13]. Although larger, the RtP is the furthest away from the synaptic terminal and unable to provide vesicles to facilitate synaptic transmission as readily as other pools [12, 14, 15]. We model this effect as a reduction in coupling between neurons. Our vesicle depletion and replenishment model is expressed by Eqs (4) and (5) (parameters in Table A in Supplementary Material), and presented in detail in Methods section *Modelling the effect of Vesicle Depletion and Replenishment on Coupling Strength*. Hence, Fig 1D exhibits a fast initial drop in coupling strength as the RRP and RP are quickly depleted, and the RtP becomes dominant.

The second key modification of the Kuramoto model that was found to be necessary for reproducing the slow ERNA dynamics is the second-order Fourier series coupling function (Fig 1E, and see Methods section *Coupling Function and PRC*). The coupling function is denoted by *f* in the model (see Eq (1)), and captures how each neuron’s phase is influenced by the phase of other neurons. The fitted coupling function has a strong second harmonic as well as a negative shift (Fig 1E). We show in Supplementary Materials Section D.1 that neither a coupling function derived from the Hodgkin-Huxley model, nor a sine coupling function are suitable to reproduce the slow ERNA dynamics. We also demonstrate in Supplementary Materials Section D.1 that the zeroth and second harmonic of the coupling function have a larger impact on the ERNA frequency decay than the first harmonic. The importance of the second harmonic is related to the clustering behavior exhibited by the model (see Section B in Supplementary Materials). In the Kuramoto model, stimulation can advance or delay a neuron’s spiking by advancing or delaying the phase of individual oscillators. This effect is captured by the phase response curve (PRC), which characterises the change in an individual oscillator’s phase as a function of the phase at the time of stimulation and is denoted by *g* in Eq (1). For this parameter set, a sine PRC (Fig 1F) was sufficient to provide an accurate fit to the features, which further limited the number of parameters that needed to be optimised.

### Long-Term Spectrogram

While Fig 1A captures the slow dynamics of ERNA over more than a minute on-stimulation, Wiest et al. provide a series of on-DBS windows and pauses that demonstrate how reproducible ERNA is [1]. Additionally, Fig 1A does not demonstrate the ability of the model to return to baseline following stimulation, a key observation of ERNA dynamics. We address this by simulating the same DBS on/off paradigm used in [1] and depicted in Fig 2A. The spectrogram (with the addition of white noise) appears similar to that of the data (compare Fig 2A and Fig 2B), and consistently returns to baseline after stimulation is turned off.

Wiest et al. outline four key characteristics of the ERNA spectrogram, see annotations in Fig 2A: (1) Initial high frequency, high amplitude response; (2) Steady state reached at a peak of approximately 250Hz after around 100 seconds of stimulation; (3) Activity off-stimulation at the low gamma/high beta frequency range of approximately 40Hz, and; (4) low amplitude, high frequency activity at around 250Hz off-stimulation [1]. The long-term spectrogram from the model (Fig 2B) demonstrates most of these characteristics and is capable of explaining all of them through model dynamics. Characteristics 1,2 and 3 can each be seen in the spectrogram regardless of the length of the DBS window. Characteristic 3 cannot be seen in first off-stimulation period as the model is in a settled state. However, in each of the following periods between DBS windows the amplitude of the low frequency activity (characteristic 3) decreases with time after DBS termination in both the data and the model. While the signal appears to remain at a high amplitude immediately after stimulation termination, the stable dynamics off-stimulation are low amplitude and low frequency in the model. Characteristic 3 is slightly more finely tuned in the model, but occurs at a comparable frequency to the data. Characteristic 4 cannot be seen in the model’s spectrogram, however, this is approximately the mean natural frequency of the oscillators in the model. Neurons occasionally being released to their natural frequency by a decrease in network coupling due to stronger fluctuating neural synchronisation or noise in the data could therefore account for characteristic 4.

### Variable Stimulation Frequency

We simulate the fitted model for various stimulation frequencies as was done in [6]. The variation in stimulation frequency is captured by the delta term in Eq 4 and will have the effect of depleting all three vesicle pools further and more quickly with higher stimulation frequency. Without making any changes to the model besides the frequency of the pulse train we are able to replicate most features of the ERNA frequency decay over time at each stimulation frequency reported in [6] (compare Fig 3A and B). All ERNA frequency decays initiate at similar high frequencies around 380Hz in both the model and the data. The decays also stabilise after around 60 seconds of stimulation in both model and data. The 180Hz stimulation frequency line finishes at around 270Hz and the 70Hz stimulation frequency line at around 310Hz, much higher than the 100Hz stimulation frequency line, in both model and data. The 180Hz line finishes higher than the 150Hz line in the data, however, in the model the 180Hz line finishes lowest. The 150Hz line finishes above the 130Hz line in the model, showing that there is enough variability in the model for crossover amongst the various stimulation frequencies. The amplitude lines also demonstrate similarities in both cases (Fig 3C and D). The 180Hz stimulation frequency line is initially higher than the other stimulation frequencies, with the 130Hz line starting above the 150Hz line in both the model and the data. Additionally, the 100Hz and 70Hz stimulation frequency lines are lower at around one in both the model and the data. Later in the stimulation window of the model, the 180Hz stimulation frequency line decreases quickly towards the 130Hz stimulation and falls below both 150 and 130Hz lines after 40 seconds on-stimulation which lines up closely with the data. Both frequency and amplitude lines of the model response have little variation regardless of the realisation of noise, natural frequency distribution and initial conditions (Fig 3B and D).

**Fig 3.**
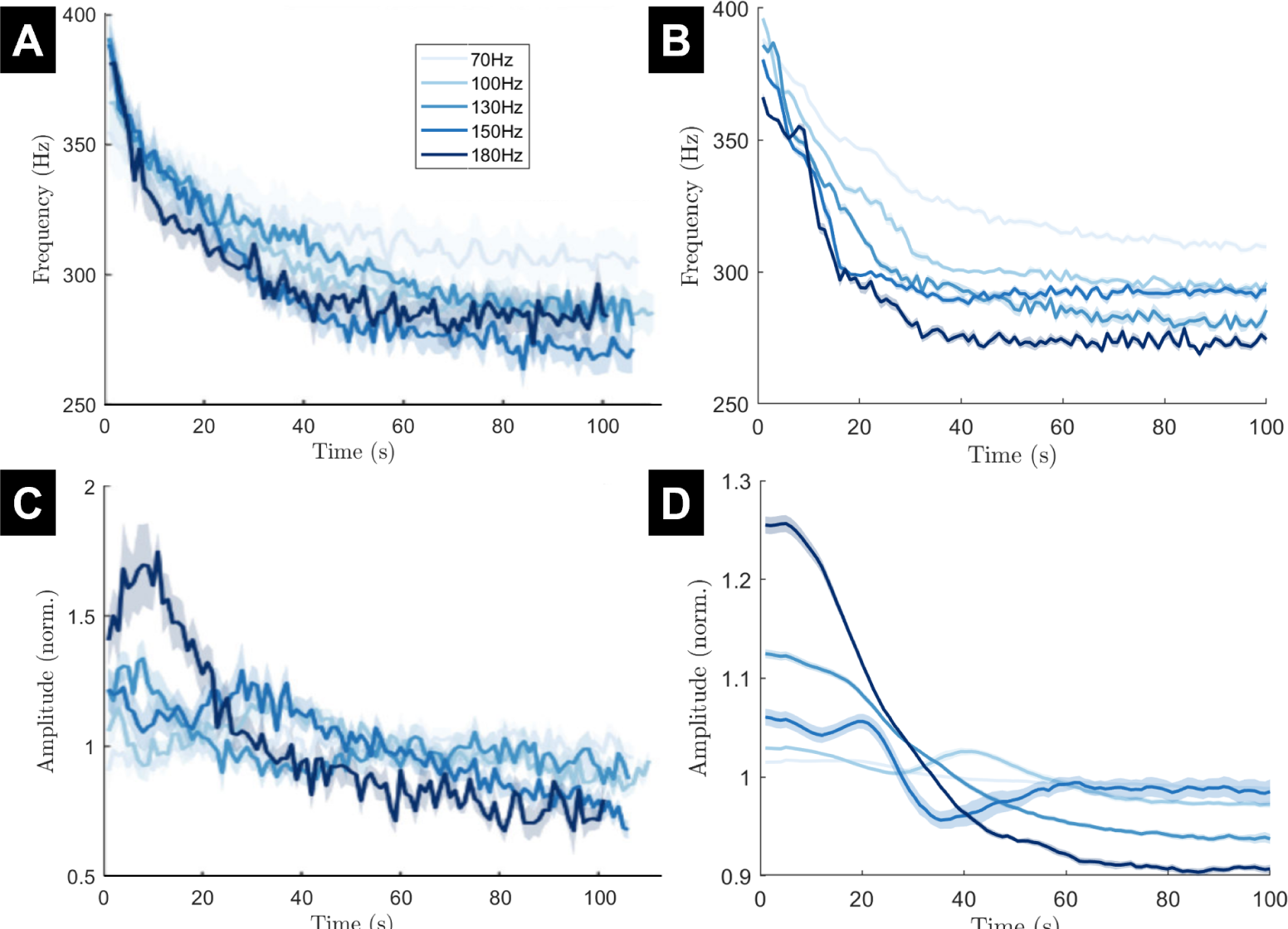
The frequency and amplitude of ERNA over time in response to various stimulation frequencies in both data and the fitted model. (A and C) The data ERNA frequency decay and amplitude variation, respectively. Stimulation amplitude was maintained at the patient’s therapeutic intensity. (B and D) The ERNA frequency decay and amplitude variation, respectively, from model simulations. The 130Hz line of the model simulation has the same parameters as the simulation from Fig 2. Mean (solid line) and standard error of the mean (faded region around solid mean) calculated over 15 trials with different realisations of noise. The colour for each stimulation frequency is the same in data and model panels. Panels A and C are adapted from [6].

### Variable Stimulation Amplitude

We also simulate the fitted model for various stimulation amplitudes as was done in [6]. Unlike stimulation frequency, the inclusion of variable stimulation amplitude needs to be made through a biophysically-motivated model parameter adjustment. Hence, we increase the probability of vesicle release with increasing stimulation amplitude (see Methods section *Variable Stimulation Amplitude*, Eq 11). This adjustment provided enough flexibility to closely reproduce the ERNA frequency decay at each stimulation amplitude (Fig 4A and B). All frequency decays initiate around 380Hz in both the model and the data. The greatest stimulation amplitude consistently produces the largest frequency decay, dropping to 300Hz after around 30 seconds before continuing below that. The lowest stimulation amplitude generally has the highest frequency, finishing around 320Hz after 30 seconds like the data and 300Hz at the end of the stimulation period. The amplitude lines of the model also demonstrate similarities to the data (Fig 4C and D). The highest stimulation amplitude initially has the highest amplitude response and is grouped with other higher stimulation amplitude lines. The lowest stimulation amplitude in the model initiates and remains around an amplitude of one, similarly to the 1mA stimulation amplitude line in the data.

**Fig 4.**
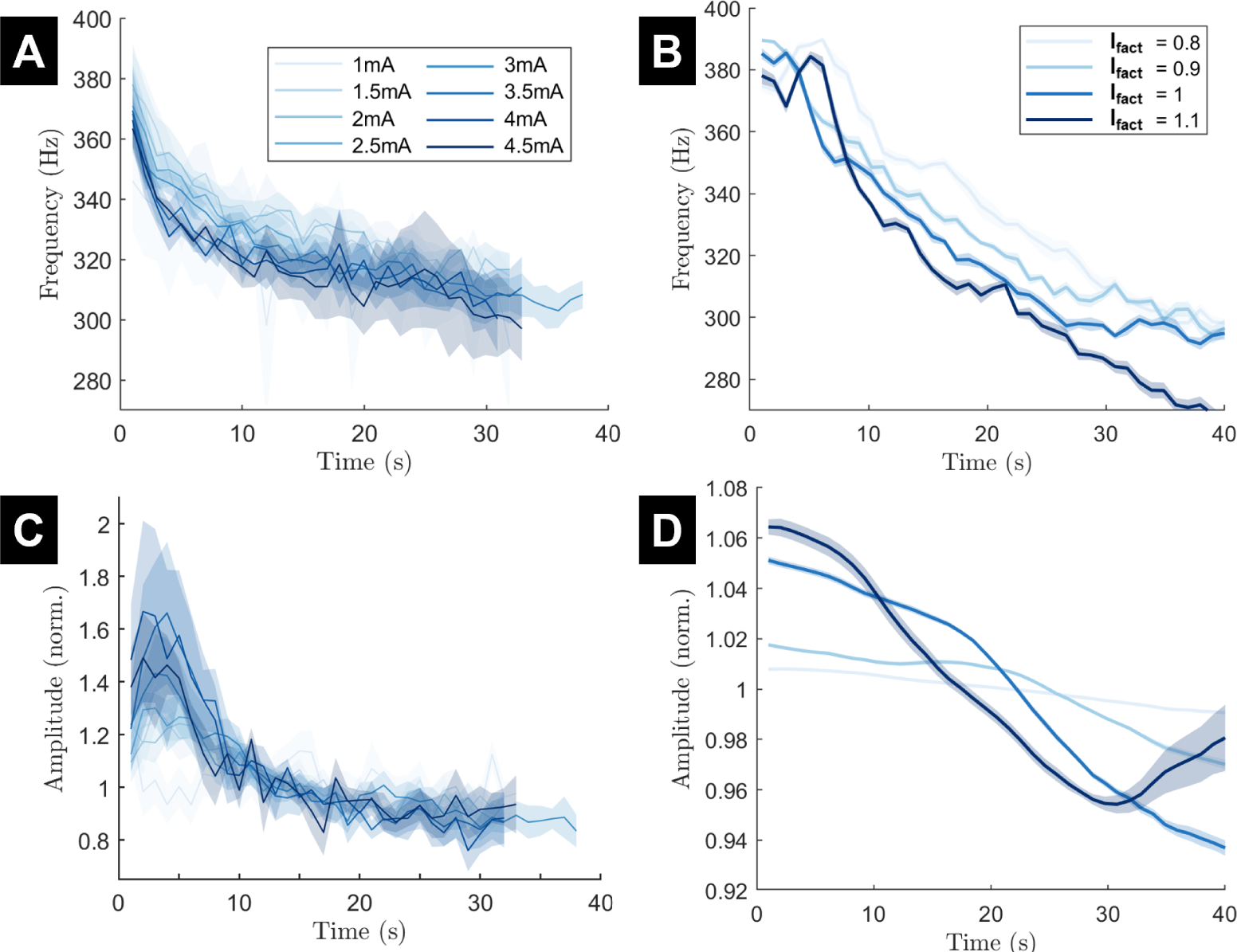
The frequency and amplitude of ERNA over time in response to various stimulation amplitudes in both data and the fitted model. (A and C) The data ERNA frequency decay and amplitude variation, respectively. Stimulation frequency is set at 130Hz. (B and D) The ERNA frequency decay and amplitude variation, respectively, from model simulations. The *I_fact_* = 1 line of the model simulation has the same parameters as the simulation from Fig 2 and approximately corresponds to 3mA. Mean (solid line) and standard error of the mean (faded region around solid mean) calculated over 15 trials with different realisations of noise. The colour for each stimulation frequency is the same in data and model panels. Panels A and C have been adapted from [6].

### Variable Medication State

We simulate the fitted model in the on- and off- medication states as was performed in [6]. We make a biophysically-motivated model parameter change to reflect medication-dependent vesicle pool changes. This change produces slower replenishment and therefore greater depletion of the RtP vesicles in the off medication state, which lowers the coupling strength during stimulation (see Methodology section *Simulations for Validation with Variable Systems - Variable Medication State*). This change provides enough flexibility to the model to effectively capture the ERNA slow dynamics on- and off-medication (Fig 5). The frequency decays of on-medication and off-medication states initiate at the same frequency of around 380Hz in both the model and the data (Fig 5A and B). The model and data frequency responses remain at similar levels and the frequency decays follow similar time constants. Medication states significantly separate in the data around 60 seconds following stimulation onset. Statistical analysis of the model output demonstrates that significant separation of the frequency responses between medication states first occurs at a similar stage (see Methodology section *Statistics* for details). This occurs after 40 seconds of stimulation in the model and is likely detected earlier in the model than the data due to the smaller standard error of the mean in both the on- and off-medication simulations. From this point theon-medication remains higher, finishing the 120 second stimulation window at around 280Hz in both model and data. The off-medication state drops to a lower frequency, stabilising around 260Hz. The two frequency response lines are significantly separated (p*<*0.05) for the remainder of the stimulation period from 67 seconds onwards in the model. The amplitude lines also demonstrate similarities, with the off-medication line remaining higher than the on-medication line early in the stimulation window. Towards the end of the stimulation period both amplitude lines drop to lower values than at the start of the stimulation window in both the model and the data (Fig 5C and D).

**Fig 5.**
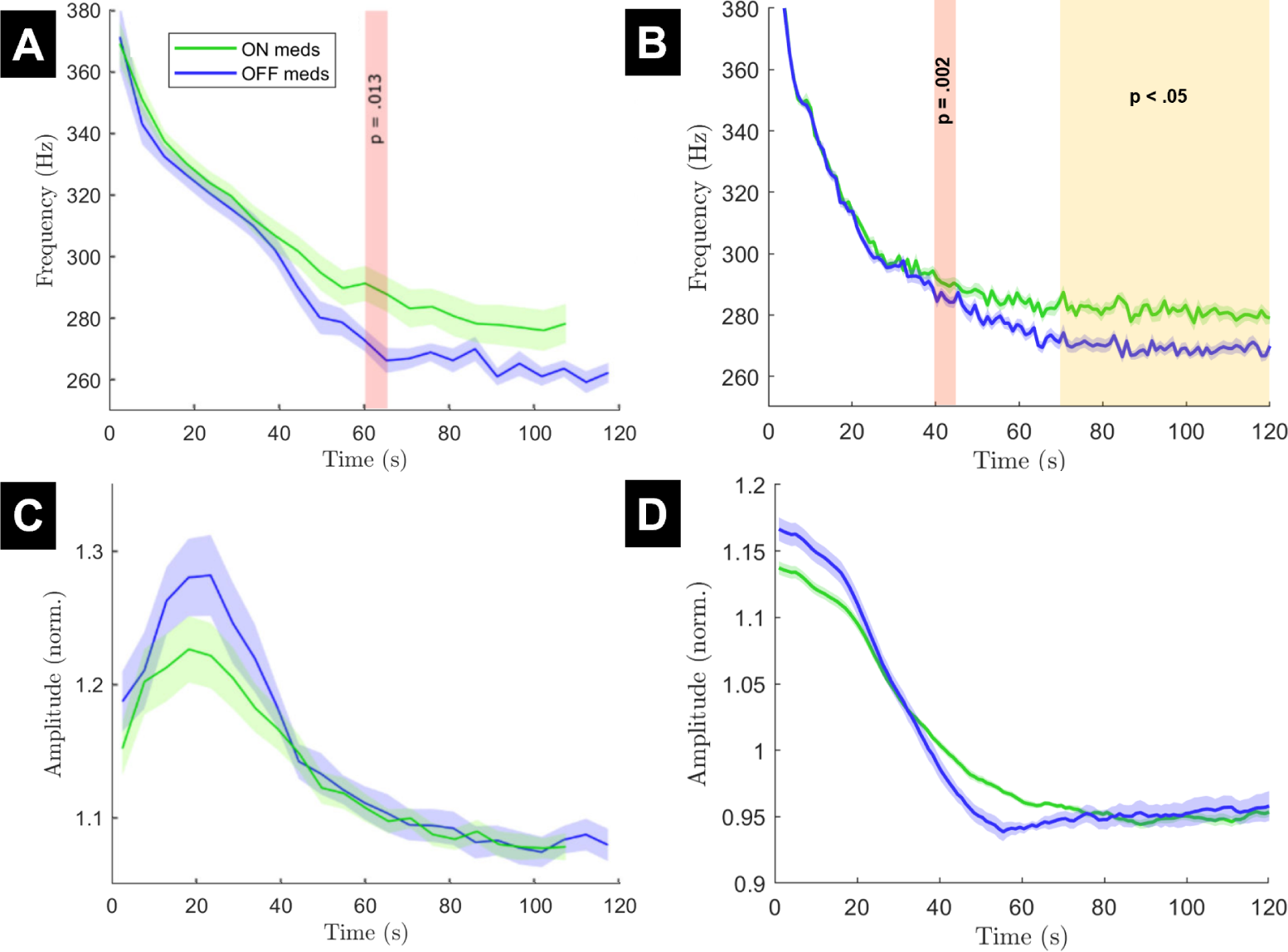
The frequency and amplitude of ERNA in response to medication state in both data and the fitted model. (A and C) The data frequency decay and amplitude variation, respectively. The red bar in panel A indicates the time after which the frequency decays of each state are determined to have significantly separated. (B and D) The frequency decay and amplitude variation, respectively, from model simulations. The on-medication line of the model simulation has the same parameters as the simulation from Fig 2. Mean (solid line) and standard error of the mean (faded region around solid mean) calculated over 15 trials with different realisations of noise. For panel B, statistical tests in line with the data were performed. We identified the points where significance was first achieved at the same level as the data (the red bar, p = 0.002), and when it was maintained for the remainder of the stimulation window (the yellow bar, p*<*0.05). The colour for each medication state is the same in the data and the model panels. Panels A and C are adapted from [6].

### Post Stimulation Bursting

The responses to post stimulation bursting (10 pulses of stimulation at 130Hz every second following termination of a long period of stimulation) also appears to show similarities between the model and the data (Fig 6). This condition captures the cumulative effect of successive, brief periods of vesicle depletion and replenishment (different time scale than in Fig 2).Vesicle depletion and replenishment over the bursting periods is described in the model by Eqs. 4 and 5 as before, and can be visualised in Fig 6E. There was no fitting performed to features specific to post stimulation bursting. In the data, the frequency response increases back up towards the early continuous stimulation state within the 20 seconds of bursting stimulation (Fig 6A). A similar increase is seen in the model, reaching approximately the same frequency at similar rates of recovery (Fig 6B). Amplitude response of ERNA also appears to increase and reach a similar level at the end of the 20 second bursting stimulation window in both the model and the data (Fig 6C-D). The dynamics of these evoked responses change at a rate determined by the vesicle replenishment and coupling strength recovery (Fig 6E). Coupling strength drops to RtP dominance during each series of stimulation bursts and recovers to RRP dominance between pulse bursts.

**Fig 6.**
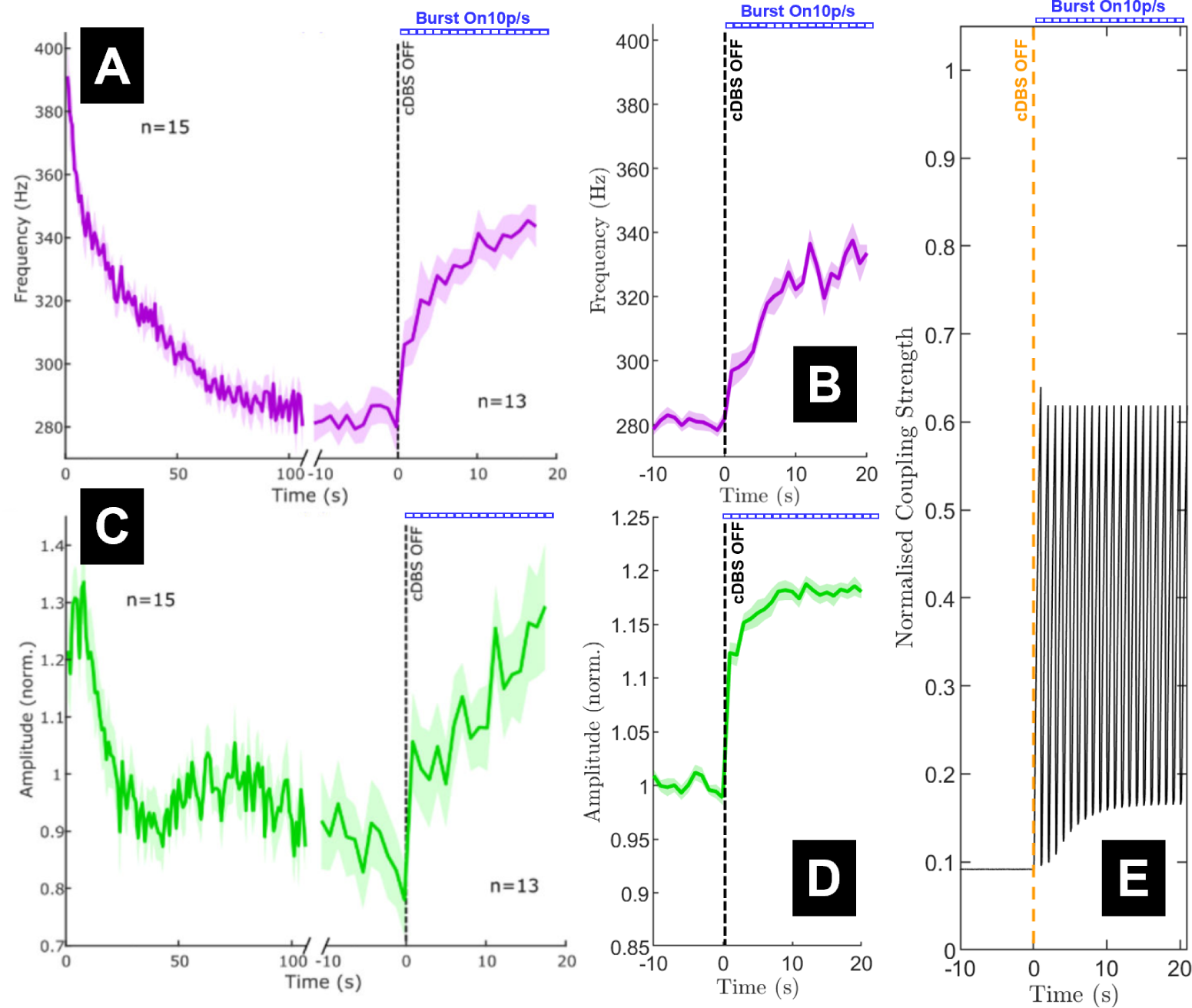
The frequency and amplitude of ERNA in response to bursting stimulation post continuous deep brain stimulation (cDBS) termination in both data and the fitted model. (A and C) The data frequency decay and amplitude variation, respectively. (B and D) The frequency decay and amplitude variation, respectively, from model simulations. Mean (solid lines) and standard error of the mean (faded region around solid mean) calculated over 15 trials with different realisations of noise. (E) Coupling strength normalised by *k_mu_* through continuous stimulation to termination at time 0 (orange dashed line) with 10 pulse bursts at 130Hz every second for the next 20 seconds. The blue dashed line in every panel (Burst On 10p/s) indicates the presence of bursting stimulation of 10 pulses at 130Hz every second following the termination of cDBS. Panels A and C are adapted from [6].

### Individual Oscillator Perturbation

While coupling between STN neurons (and how it varies over stimulation) is crucial in our model’s ability to reproduce ERNA dynamics, results presented in [9] portray the STN as a mostly disconnected network of neurons. Steiner et al. showed that stimulating individual STN neurons to the point of firing does not cause neighbouring STN neurons to fire. We therefore simulate perturbations to single oscillators in the network in order to determine whether these results can be replicated in the model while still having connectivity between STN neurons. We individually perturb seven of the oscillators from our network (all-to-all connected) with four pulses at 130Hz and the same stimulation amplitude as the model was fitted to (3mA). In [9], four pulses at 20Hz were used, in this study we proceed with 130Hz as the model did not capture the long term 20Hz response (see Supplementary Materials Section C.2). This is a conservative choice, since 130Hz stimulation elicits faster phase progression (i.e. progression towards spiking) of the oscillators in the model. We record the rate of phase progression in all seven oscillators with each perturbation and compare this against the phase progression of the oscillators in the absence of stimulation to any part of the network. There appears to be no marked change in phase progression rate of the oscillators across the network in response to a single oscillator being perturbed (Fig 7). All of the oscillators not being directly stimulated respond similarly showing a small increase in phase progression, but not enough to experience any additional firing (*<* 2*π*, corresponding to one cycle). This is seen across the network regardless of the oscillator being perturbed or observed.

**Fig 7.**
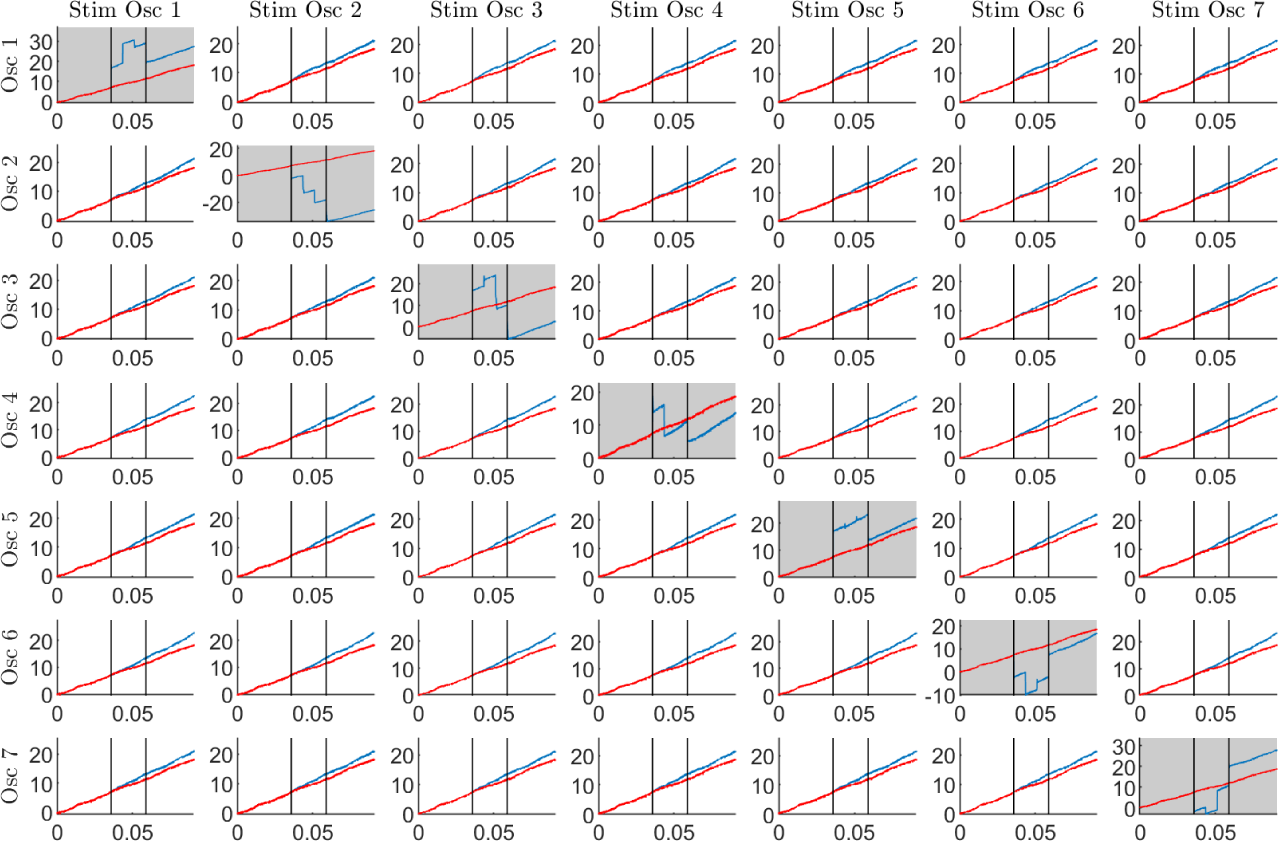
The phase progression of individual oscillators in response to the perturbation of a single oscillator in the network for comparison against data [9]. Observing phase progression of oscillator indicated by row title during stimulation of oscillator indicated by column title. The grey panels represent a case where we are recording the phase progression from the same oscillator as is being stimulated. White panels indicate a recording from an oscillator not receiving micro-electrode equivalent stimulation. The red lines on each plot represent the phase progression of each oscillator (number indicated by row heading) in the absence of stimulation to any oscillator in the network. The blue lines represent the phase progression of the oscillator in response to stimulation of a single oscillator. Stimulation is applied at 130Hz for four pulses at the fitted stimulation amplitude between the two vertical black lines as was done in the data [9].

### Replication of In Vitro Slice Experiments

The behaviour of mostly disconnected oscillators in the model matches data from in vitro slice recordings of the rat STN (Fig 8). Rat STN neurons in slices were shown to fire around 20Hz off-stimulation [16, 17]. In the presence of constant current electrical stimulation, firing rate increases rapidly to over 200Hz at low stimulation amplitude (*<*0.5nA) [16, 17], up to over 500Hz in some cases [18]. This was consistent with high-frequency pulsatile stimulation (*>*120Hz, 200*µ*A) where firing rate of individual STN neurons was seen to increase to over 200Hz [19]. Using two oscillators to capture the mostly disconnected network, we incorporate the absence of GABAergic input from the GPe in slices by increasing the shift of the coupling function (the noughth order term, *f*_0_, of the Fourier series presented in Eq.6) such that the oscillators fire at around 20Hz off-stimulation (Fig 8A). In the presence of constant current stimulation, the two oscillators progress at a frequency greater than 500Hz immediately following stimulation onset (Fig 8B). We used a stimulation amplitude scaled to match the intensity of pulsatile stimulation from the parameter fitting (see Methodology section *Replication of In Vitro Slice Experiments* for details). This was likely a higher stimulation amplitude than these slice experiments used, as the model was fitted to pulsatile stimulation data of around 3mA. Despite the likely disparity in the amplitude of stimulation, the firing rate on-stimulation matches the observations of neuronal firing rate increasing to over 500Hz with stimulation [18]. When vesicle depletion is included in this network, the approximate frequency of the oscillator firing rate stabilises around 300Hz.

**Fig 8.**
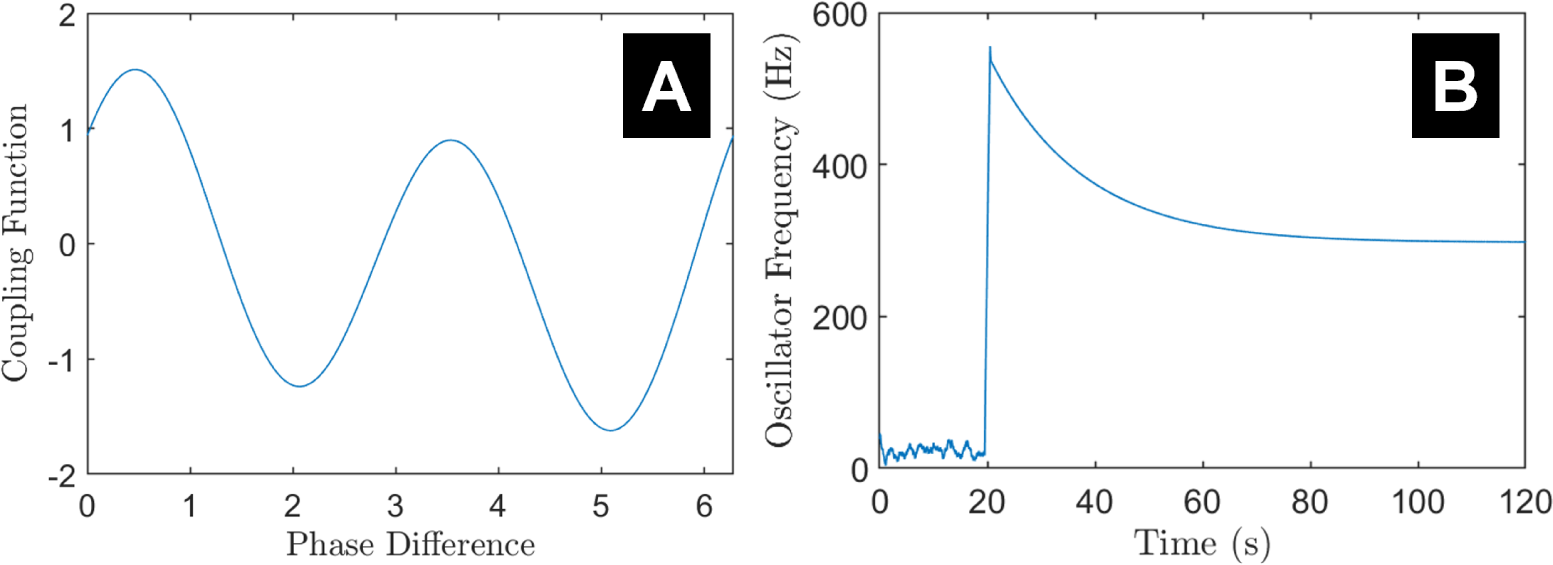
The approximate frequency of individual oscillators in a two oscillator network with the adjusted coupling function and constant current stimulation for comparison of mostly disconnected in vitro slice experiments. (A) Adjusted coupling function after changing noughth order coefficient. (B) Individual oscillator frequency in two oscillator network over constant stimulation current injection window of 100s following onset after 20 seconds off-stimulation.

## Methods

In this section, we first provide an overview of the computational model, starting with an introduction to the Kuramoto model as well as the details of the modifications to the model, in particular the effect of vesicle depletion and replenishment on coupling strength. We then present the fitting process used to find model parameters that reproduce key characteristics of the slow dynamics of ERNA. Lastly, we describe methodological details for all simulations performed in this study based on different experimental paradigms.

### The Kuramoto Model

Our approach is to attempt to model the ERNA using as simple a model as possible. We therefore consider the Kuramoto model, which has been frequently used to represent oscillatory neural activity as well as the effect of DBS [20–25], and is one of the simplest models that can describe populations of interconnected neurons. The basic Kuramoto model describes a series of interconnected phase oscillators [26], in our case representing neurons [22, 27]. The evolution of the *i^th^* oscillator’s phase *θ_i_* is given by the stochastic differential equation

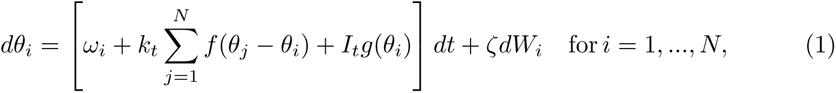

where *ω_i_* represents the natural frequency of the *i^th^* oscillator. The (homogeneous) coupling strength between oscillators at time *t* is *k_t_*, and *f* represents the coupling function, which is a 2*π*-periodic function of phase differences. The phase response of individual oscillators is described by the 2*π*-periodic PRC which is denoted *g*, and *I_t_* is the intensity of stimulation at time *t* modelled as

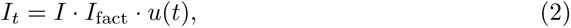

where *I* is the fitted stimulation amplitude and *I*_fact_ is a factor included to vary stimulation amplitude. The stimulation pulse train *u* at time point *t_i_* represents a simplified DBS pulse train obtained as

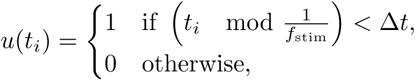

where *f*_stim_ is the stimulation frequency and Δ*t* the simulation time step. We are assuming that all oscillators perceive the same level of stimulation. Intrinsic noise is added to each oscillator through independent Wiener processes *W_i_*. The noise standard deviation is set by *ζ*. The number of Kuramoto oscillators in the model is given by N, which is chosen to be 50 for this study unless stated otherwise. To measure synchrony in the network, we use the order parameter given by

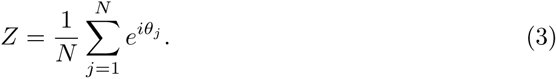

It is necessary to introduce a series of modifications to the Kuramoto model in order to incorporate dynamics that can adequately capture the ERNA response. While we do not include delays or heterogeneous coupling in this study, the modifications described below provide enough versatility in the model dynamics to capture the nonlinear behaviour of ERNA.

#### Modelling the effect of Vesicle Depletion and Replenishment on Coupling Strength

Vesicle depletion and progression towards synaptic failure has been suggested as an underlying mechanism of the ERNA frequency decay [6]. However, no clear mechanism describing how vesicle depletion may influence the dynamics of ERNA has been proposed. We will develop the vesicle depletion hypothesis in this study and sketch how it could explain the underlying dynamics of ERNA.

Rizzoli and colleagues proposed the three vesicle pool model of synaptic transmission (see Fig 9A) from the observation of three stages of fluorescence in goldfish bipolar cells [12]. Firstly, the readily-releasable pool (RRP) is a small but easily-mobilised group of vesicles. Due to their proximity to the synaptic terminal and small number, RRP vesicles are often depleted within a second of stimulation onset and recycle within seconds. Secondly, the recycling pool (RP) is a slightly larger pool of vesicles which are less easly mobilised, but still deplete and recycle within seconds. Finally the reserve or resting pool (RtP) represents the majority of vesicles at the synaptic terminal. The RtP is the least easily mobilised vesicle pool and its vesicles are depleted over minutes (depletion only occurs under strong stimulation) and recycle over a similarly long time period. The vesicle pools are recruited in this sequential order. The RRP is the primary vesicle pool responsible for synaptic transmission under physiological firing conditions. It is able to recycle vesicles between action potentials under physiological conditions, and was first observed as quanta docked to the synaptic terminal [28].

**Fig 9.**
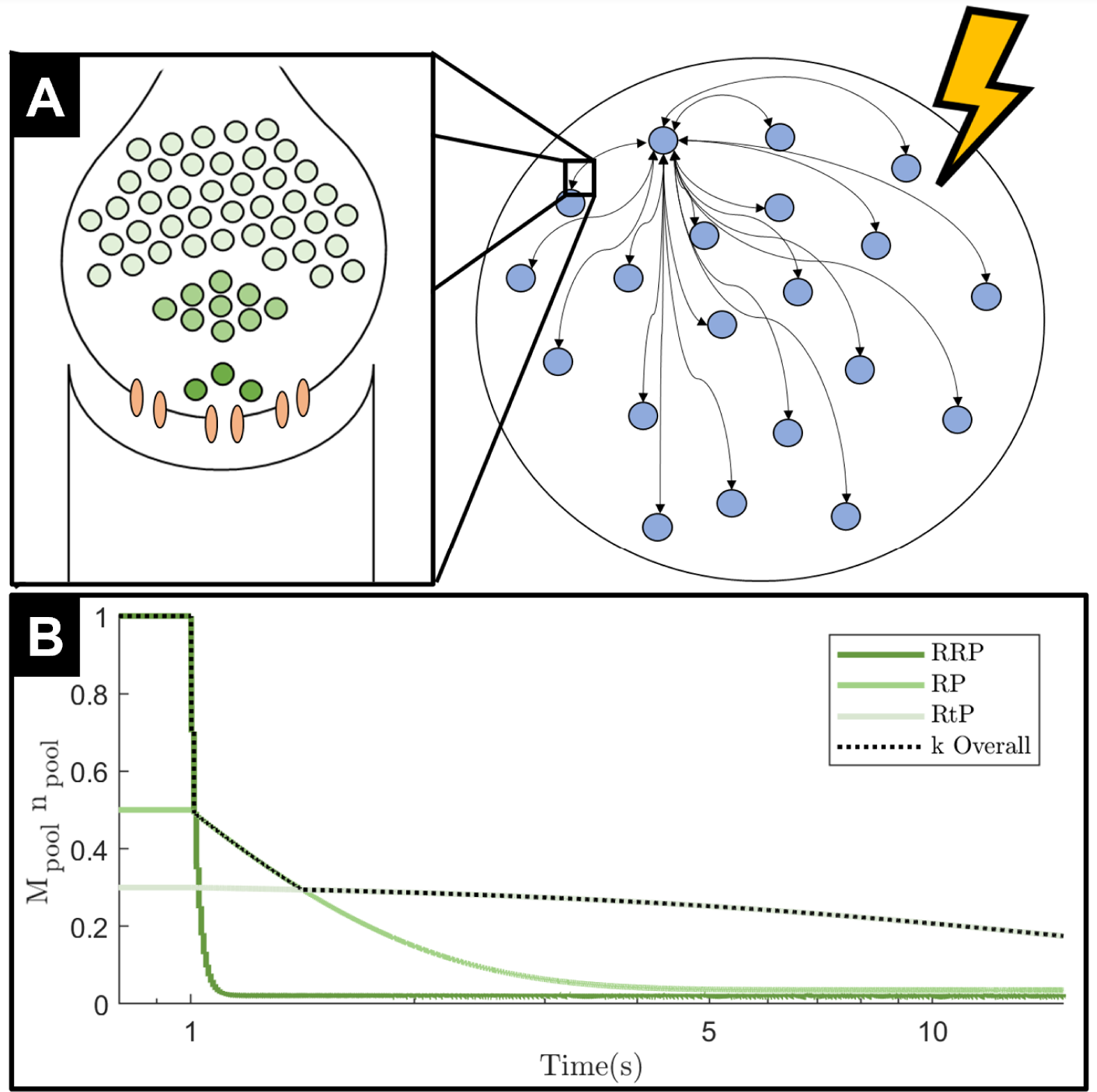
Modified Kuramoto model. (A) Right: One population of all-to-all connected Kuramoto oscillators (blue circles), which all receive stimulation of the same magnitude. Only showing bi-directional connections to one example oscillator. Left insert: We zoom in on the synaptic connections between each oscillator which are affected by vesicle depletion. The closest vesicle pool to the synaptic terminal is the RRP (darkest green), followed by the RP (lighter green). The pool furthest away from the synaptic terminal is the RtP (lightest green). Vesicle pool colours are consistent with colours used in panel B. (B) Coupling strength decay curve, with stimulation starting at 1s. The three green colours represent the contribution of each of the three vesicle pools. The black dotted line is the output of the maximum function in Eq 5, which dictates the overall coupling strength.

In the context of ERNA, however, the STN is experiencing unphysiological stimulation above the threshold required to observe vesicle depletion [11]. Under these conditions, simulations show the RRP depleted within approximately 10 pulses of stimulation at 100Hz [13]. Hence, after prolonged exposure to unphysiological stimulation the majority of the vesicles responsible for synaptic transmission will belong to the RtP [14, 15]. During the early stages of stimulation and vesicle depletion, the number of vesicles remaining at the terminal appears to follow an exponential decay [15]. Vesicle depletion of STN axonal efferents has been observed in response to STN-DBS in both in vivo and in vitro recordings [29].

Computational models characterising the effect of vesicle depletion on synaptic connectivity and the response of a vesicle pool to stimulation have previously been developed [30, 31]. In the context of the three vesicle pool model, the interaction of multiple vesicle pools has also been investigated across a range of pool occupancies [32, 33]. We employ the first order kinetic model used in [30] to track vesicle pool occupancy. Given the time scale differences over which each pool depletes and replenishes, we model the dynamics of each pool independently. This simplified approach is sufficient to capture the three different time constants of each vesicle pool (see Fig 9B) as seen in data [12]. Following [30], we model the occupancy of each vesicle pool *n*_pool_ ∈ [0, 1] (with pool ∈ {RRP, RP, RtP}) as

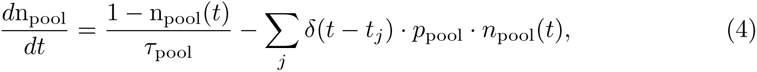

where *τ*_pool_ is the time constant of replenishment and *p*_pool_ is the probability of vesicle release in response to a stimulation pulse for the fitted stimulation amplitude for each vesicle pool. The values used for these parameters are estimated from the time taken to deplete and replenish each of the vesicle pools, respectively, according to data [12, 13] – see Section *Fitted Model Parameter Set* in Supplementary Material. In modelling the evolution of pool occupancy, we focus on the effect of stimulation and ignore spikes not triggered by stimulation. Thus, *δ*(*t − t_j_*) is a Dirac function where the *t_j_*’s correspond to the time points of stimulation.

We model the effect of depletion and replenishment of the three vesicle pools (given by Eq (4)) on coupling strength based on three assumptions. First, activity at a synaptic terminal directly corresponds to the number of vesicles released [34], and if fewer vesicles are released into the synapse, the effect of pre-synaptic activity on downstream neurons will be weaker. Thus, we model the effect of vesicle dynamics on coupling strength as proportional to the number of vesicles available.

Second, we translate the occupancy of each vesicle pool into its effect on coupling strength using a factor *M*_pool_, which we estimate for each vesicle pool using data as the product of both the pool’s relative size [12] and the inverse of its approximate distance from the synaptic terminal [12, 35–40] (less readily-releasable vesicles further from the synaptic terminal have a lower release probability for a given stimulation [35]). The values used for the pool-coupling factors, *M*_pool_’s, are given and justified in Section *Fitted Model Parameter Set* in Supplementary Material.

Third, at any point in time, we only consider the effect on coupling strength of the dominant pool (largest *M*_pool_ *n*_pool_(*t*) at time *t*). This reflects the sequential nature of vesicle pool depletion and how one pool tends to dominate synaptic transmission at any point in time depending on depletion levels. It also allows the vesicle depletion curve to take the form of an exponential decay with three time constants (Fig 9B) as seen in [12]. Moreover, while on-stimulation coupling strength is impacted (in succession) by the depletion of each vesicle pool, the recovery of coupling strength following stimulation termination is chiefly determined by the replenishment of the RRP. Other vesicle pools do not appear to play a significant role as synaptic activity recovers at a similar rate regardless of the stimulation parameters used during the stimulation window [11] (i.e. regardless of how depleted the other pools are), which is consistent with this third assumption. Since the RRP is replenished much faster than other pools (within seconds following stimulation termination [12]), the RRP quickly resumes being the primary vesicle pool for synaptic transmission.

Based on these three assumptions, the coupling strength at time *t* is obtained as

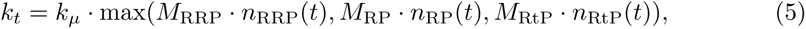

where *k_µ_* is the steady state coupling strength of the network off stimulation.

#### Coupling Function and PRC

In addition to the modulation of coupling strength over time when the network is exposed to stimulation, we incorporate a second-order Fourier series coupling function *f* into the model and fit its coefficients (*f_i_*), *i* ∈ {0, .., 4 to data. The coupling function is thus given by

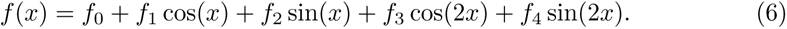

Compared to a sinusoidal or Hodgkin-Huxley-derived coupling function (explored in Supplementary Materials Section D.1), this more general coupling function allows the model to capture more complex interactions between neurons.

For the majority of this work, we consider a sinusoidal PRC, i.e. *g*(*x*) = sin(*x*). The inclusion of a second-order PRC was also considered, as well as sinusoidal and a Hodgkin-Huxley PRC (see Supplementary Materials Section D.2). The form of the PRC was not as crucial to the intricate dynamics of ERNA as the coupling function was observed to be.

#### Distributions of Oscillators’ Natural Frequency

Since the STN is a nonuniform neuronal structure, oscillator’s natural frequencies are sampled from a normal distribution, *ω_i ~_ N*(Ω*, σ*^2^). We do not constrain the mean angular frequency Ω and standard deviation *σ* to a pre-determined narrow frequency band in the fitting process to provide added versatility to the model dynamics. Some in vivo and in vitro recordings of STN neurons showed firing rates of around 20Hz off stimulation [16, 17]. However, these recordings are from networks of neurons and may not represent isolated neurons’ intrinsic firing rates. This may be supported by the observation of some high-frequency activity off stimulation in the STN (see 4 in Fig 2A). We present simulations with natural frequencies constrained to 20Hz in Supplementary Materials Section E.

### Fitting Process

In order to find a set of parameters for the modified Kuramoto model that yield dynamics similar to ERNA, we fit the model to features of the slow dynamics of ERNA. The fitting methodology used in this study is similar to other computational studies [41–43] and uses the patternsearch MATLAB function to minimise a cost function representing the distance between the data features and the same features calculated in the model. The Kuramoto model was simulated using a Euler-Maruyama scheme with a timestep Δ*t* = 10*^−^*^4^.

The cost function used in the optimisation process is calculated from the on-stimulation ERNA frequency decay (slow dynamics, Fig 1B). An approximation of the peak frequency of the ERNA decay in the STN for a trial of 130Hz and 3mA STN DBS (patient on medication) was obtained using Fig 1B (taken from [1]) as

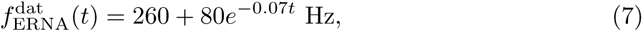

where *t* is the time elapsed since DBS was turned on. The estimate 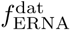 is represented by the black line in Fig 1C. The Kuramoto model was then simulated for a period of 100 seconds on stimulation following a 20 second settling period off stimulation. The model’s frequency decay 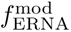 was calculated as the maximum PSD frequency between 240-400Hz for a series of one 100ms windows within the 100 second stimulation period. The cost of this feature was obtained as

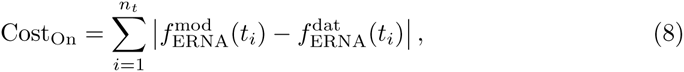

where the *t_i_*’s correspond to sampling every 100 ms and *n_t_* is the number of time samples.

Another spectrogram feature was used to limit high frequency activity before stimulation. This was done by calculating the maximum over time off stimulation of the complex modulus of the order parameter, |*Z*|_max_, a metric for the synchronisation of the oscillators in the network. The off-stimulation cost was defined as

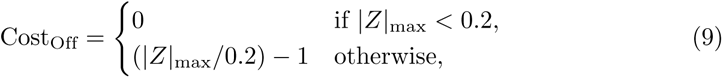

where the value 0.2 acts as a threshold under which the off-stimulation cost is zero, and above which the off-stimulation cost grows linearly with |*Z*|_max_. The threshold of 0.2 provided enough of a contrast compared to on-stimulation activity, which tended to be much closer to 1.

The on-stimulation feature was chosen as the most important to replicate, and considering the scale of both features, the total cost corresponding to the slow dynamics of ERNA was evaluated as

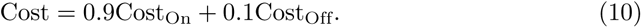

### Simulation of Various Stimulation Paradigms

To ascertain the computational model’s suitability for investigating ERNA, we validate its performance by subjecting it to various stimulation paradigms that replicate conditions explored across a range of clinical studies.

#### Long-Term Spectrogram

We replicated the long-term spectrogram reported in [1], where DBS is turned on and off in a series of periods of varying lengths for a total of 1200 seconds of continuous observations (Fig 2A). In our simulations, stimulation is initially turned on for three 200-second periods separated by 40-second off-stimulation intervals. This is followed by an 80-second off-stimulation period and five 15-second on-stimulation periods with

55-second off-stimulation intervals in between. The on- and off-stimulation period durations were chosen to align most closely with the data (Fig 2A). All stimulation periods are at 130Hz and at the same stimulation amplitude as the model was fitted to (3mA). The simulation is initialised with all vesicle pools operating at 100% occupancy. White noise is added to the order parameter (extrinsic noise) using the ‘awgn’ function of MATLAB with a signal-to-noise ratio of 20. The signal-to-noise ratio and colourbar limits were chosen by visual inspection to resemble the background activity of the data.

#### Simulations for Validation with Variable Neuromodulation

In order to replicate ERNA dynamics in the model under various stimulation amplitudes, as well as in the absence of medication, adjustments to the model have to be made. We provide simple, biophysically-motivated changes to the model and fit the changes by manual parameter tuning to the ERNA frequency in the data [6] at the end of the stimulation window. To validate these changes, we then compare the amplitude of ERNA in the model to that of the data.

##### Variable Stimulation Amplitude

Unlike stimulation frequency, the scale of stimulation amplitude in the model cannot be directly compared to the stimulation amplitude used in the data. This makes interpretation of the stimulation amplitude range more difficult. We assume that increasing stimulation amplitude will increase the rate of vesicle depletion by depleting more vesicles in response to each stimulation pulse, which corresponds to an increase in *p*_pool_. Increasing stimulation amplitude should therefore result in similar effects to increasing stimulation frequency in data [44]. We model the effect of changing stimulation amplitude by replacing *p*_pool_ in Eq (4) by

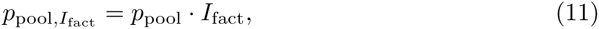

where *I*_fact_ is the stimulation amplitude modulation factor (see Eg (2)). This linear model is only appropriate for *I*_fact_ close to one. In Fig 4, *I*_fact_ is between 0.8 and 1.1 (with *I*_t_ given by Eq (2)).

##### Variable Medication State

While medication state is not a direct input to the Kuramoto model, its influence can be approximated in the model based on biophysically motivated assumptions. L-DOPA has been reported to increase long-term potentiation and depression [45]. Using the framework of spike-timing-dependent plasticity, synaptic terminals are required to maintain higher vesicle pool occupancy such that post-synaptic spikes closely follow pre-synaptic spikes (in the potentiation case). We assume that this reflects an increased rate of long-term vesicle recycling in the on-medication state. Hence, we increase the off-medication state time constant of vesicle recycling for the RtP (*τ*_OffMeds,RtP_ = 60, from *τ*_OnMeds,RtP_ = 50) by manual fitting to the final ERNA frequency after 120 seconds of stimulation.

### Replication of In Vitro Slice Experiments

We aim to reproduce firing rates of STN neurons explanted from rats exceeding 250Hz with constant current stimulation [16–18]. These in vitro slices represent neurons mostly disconnected from the rest of the STN. To replicate this condition in the model we reduce the network down to the fewest number of oscillators for which coupling can exist, two oscillators. In order to capture the absence of GABAergic input to the slice after removal from the brain, we increase the shift of the coupling function between the two oscillators. We adjust the shift of the coupling function so that both oscillators progress with a frequency of approximately 20Hz (*f*_0_ from −0.780 to −0.110 in Eq (6)) as seen in the off-stimulation data [3, 16, 18]. Constant direct current stimulation is then applied. The stimulation amplitude is adjusted from the value the full model was fitted to by a factor 130 x Δ*t* = 0.013 to deliver the same stimulation intensity per oscillator in this simulation as in the 130Hz pulsatile stimulation case in the full model. We approximate the instantaneous frequency of the *i^th^* oscillator at time *t* as

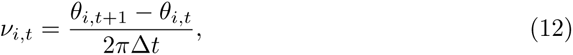

which is then smoothed (moving average, 100ms window) to reduce the effect of noise.

### Evaluation of ERNA Frequency and Amplitude over time

We visualise the computational output in a manner consistent with the approaches used in the corresponding experimental paradigms [6]. We consider the real part of the order parameter *Z* to approximate the neuronal signals from data, in line with [22]. Spectrograms are calculated both on and off stimulation using the ‘pwelch’ MATLAB function applied to the real part of *Z* over one second windows with a 25% overlap for Fig 2 (in line with [1]) and 100ms windows with a 50% overlap everywhere else as these are shorter simulations. There are no notch filters in place at stimulation frequency or the harmonics as the artefacts of stimulation in the real part of *Z* of our model are relatively small or nonexistent. The ERNA frequency evolution over time is calculated by finding the frequency of maximum PSD on stimulation in one second windows. ERNA amplitude is calculated by taking the complex modulus of *Z*. In the data, each second a pulse is skipped to allow for the frequency and amplitude of the evoked response to be recorded. While the model does not suffer from the stimulation artefact obscuring the evoked response, we take a similar approach and average both ERNA frequency and amplitude over moving windows of one second to match the windowing used in the data.

### Statistics for ERNA Amplitude On and Off Medication

We performed a statistical analysis to determine whether the frequency response differed significantly between medication states, taking the same approach as in the experimental study [6]. We used a non-parametric cluster-based permutation test with 1,000 permutations at each time point across the 15 different simulations for each medication state. We identify the stage of simulation where p-value drops below 0.05 for the first time, as well as for the rest of the stimulation window.

## Discussion

In this study, we investigated the ability of a single-population phase oscillator model with no network delays to reproduce the key characteristics of ERNA slow dynamics. We showed that when fitted to an approximation of the ERNA frequency decay obtained from patient data, the model is able to reproduce the main characteristics of ERNA slow dynamics. Key to this ability is the inclusion of synaptic vesicle depletion as a result of high-frequency stimulation. The model represents a simplification compared to models previously used to study ERNA, while also being able to reproduce characteristics over longer time periods and in response to a wide range of neuromodulatory perturbations. Importantly, the model demonstrates that ERNA slow dynamics can be reproduced with a single neuronal population without dynamic inputs from other structures. This work highlights the fundamental properties required of a phase-oscillator network to reproduce the slow dynamics of ERNA. This information may prove useful in the development of the clinical understanding of ERNA.

### Hypothesised Mechanisms and Models of ERNA

Several mechanisms have been hypothesised to explain ERNA in STN-DBS (see [6] for a summary). One hypothesis that has gained support is the network between the STN and GPe recurrently modulating activity in both structures. This recurrent network adds delays which through an excitatory-inhibitory loop would explain the promotion of oscillatory activity seen in ERNA [46]. This theory is supported by the observations that STN-DBS tends to increase firing rate in the GPe with a pattern resembling ERNA [47, 48], as well as pallidal-DBS producing ERNA in the pallidum of patients with PD [3]. It was also demonstrated that a computational model assembled in line with this theory was able to produce ERNA-type responses over an approximately 20ms time period following a stimulation pulse [2]. However, (as discussed in [6]) extended high-frequency stimulation to the STN would likely lead to depletion of the synapses connecting the STN and GPe preventing the oscillations from continuing over an extended period of time (functional disconnection) [49]. This would not allow for the persistent high frequency and high amplitude ERNA remaining after multiple minutes of stimulation observed in data. To address this shortcoming of the patterned STN-GPe network firing hypothesis, it was proposed that DBS may initiate orthodromic activation of terminals to the stimulated structure [46]. This activation could also lead to delayed excitation and inhibition, therefore promoting the oscillatory behaviour of ERNA over a longer period of time.

A recent modelling study was able to replicate some of the key ERNA characteristics over a time period of approximately 200ms using a phase oscillator network [50]. The Kuramoto network included delays between distinct sub-populations of oscillators, which would likely be a fundamental reason as to why the network was able to reproduce these evoked oscillatory responses. However, each of these sub-populations were chosen to represent separate basal ganglia structures (globus pallidus internus, ventral intermediate nucleus of the thalamus, and the STN). Hence, this model encounters the same functional disconnection contention as proposed by Bergman et al. [49]. The network could instead be chosen to represent multiple sub-populations of neurons within a single basal ganglia structure (e.g. the STN for STN-DBS), however, this would likely effect the time constants of inter-sub-population delays due to proximity and connectivity. Additionally, it has been demonstrated that there exists a diverse range of responses to stimulation within neural structures as DBS leads pass through both the STN [51] and pallidum [52]. Hence, it is important to explore how closely a single population can reproduce characteristics of ERNA to avoid these contentions, as we investigated in this study. However, when considering STN-DBS, the existence of functional connectivity in the STN has been disputed [9]. We explored the necessity of intrinsic connectivity of the single population model in *Individual Oscillator Perturbation* in the Results section.

Other computational models have looked at responses evoked by DBS [53, 54]. These studies closely reproduce characteristics of ERNA fast dynamics for approximately 0.5 to 2 seconds following the immediate initialisation of stimulation. They achieve this through including multiple structures with delays. We instead focus on reproducing long-term ERNA dynamics over multiple minutes, which had yet to be explored. Additionally, we further expand on the work of previous models by focusing on a model of a single structure, the STN. This avoids the reliance on delays from the network to produce the resonant activity, which would likely experience synaptic failure over this longer time scale [49]. This has not been previously considered in models due to a focus on short-term responses. In fact we build synaptic vesicle depletion into the model, which is a fundamental reason the model is capable of reproducing the long-term ERNA dynamics. Additionally, by focusing on a single structure we reduce the number of model parameters which will help to prevent overfitting the model to data. We go beyond these previous studies by modelling ERNA over longer periods of stimulation, and in response to a wider variety of stimulation paradigms to validate our approach.

### Proposed Computational Model

Across the many ERNA characteristics discussed in this study, we provide a preliminary validation for the modified Kuramoto model across multiple states and inputs. The model demonstrates that it has the required versatility to replicate the frequency shift between off and on-stimulation states as well as the slow dynamics of the spectrogram. The model provides a close reproduction of the frequency decay, despite being a simplification of the neuronal system that it is representing. Furthermore, we attempt to biophysically ground and validate the model.

#### Validation with Variable Neuromodulation

The model is able to replicate the frequency and amplitude behaviours for variable inputs and states. In some cases, biophysically-motivated parameter changes are made by manual parameter tuning to match the initial and final frequency values in each of these conditions. However, these parameter changes also provide amplitude responses similar to the data without using these responses as features for fitting. The parameter tuning is restricted to one parameter and only by a small factor, which does not change the fundamental model dynamics. No tuning was required for the variable stimulation frequency example which provided the closest replication of the data for both the frequency and amplitude responses.

All variable neuromodulation approaches provided comparable approximations to ERNA dynamics when comparing the mean of the model simulations (Figs 3, 4 and 5). However, the standard error of the mean is much smaller in the model than the data. Each model simulation corresponds to different realisations of natural frequencies, initial conditions and noise. In the data the different recordings are from different subjects. Hence, to reproduce the variability observed across different subjects we would likely be required to change the parameter set for each realisation.

#### Importance of Multiple Vesicle Pools

The key modification of the computational model that enables the replication of long-term ERNA characteristics is the introduction of synaptic vesicle dynamics with multiple vesicle pools. The results (Figs 1, 3, 4 and 5) appear to demonstrate frequency decays with at least two rates, in addition to the very fast RRP dynamics. We interpret these different rates of frequency decay as the initial depletion of the RP within the first 5 seconds, followed by depletion of the RtP in the remainder of the stimulation window (see Fig 9B). The RRP plays a crucial role in determining the frequency of the network immediately before and after stimulation onset as well as in the post stimulation recovery.

#### Considerations for Other Neural Structures and Across Patients

Recent publications have shown that ERNA is not solely a property of the STN [3], however the frequency response can be quite different between structures in particular. While maintaining a similar coupling function, the interaction between individual oscillator natural frequencies as well as coupling strength in the network can be modified to accommodate these changes. In the off-stimulation state, only natural frequency and coupling strength affect the phase progression of each oscillator. These parameters and how the coupling strength evolves with time on stimulation (through parameters *τ*_pool_ and *p*_pool_) can be changed to control the frequency that ERNA appears at and decays to. In structures where ERNA is not observed (for instance in the visual cortex [5]) vesicle depletion is still expected to be present. We speculate that the other key properties of the model are the likely factors preventing these structures from producing ERNA (i.e. not having the high natural frequency of the neurons and a coupling function similar to Fig 1E).

### Model Predictions for Further Validation

This study has compared model predictions to previously published ERNA data, and in this section we highlight predictions corresponding to conditions for which experimental ERNA data is not available at the time of writing.

#### Only One Neuronal Population’s Dynamics are Required to Produce ERNA

The model used in this study only requires one neuronal structure with no reciprocal feedback between different structures. Long-term recordings from the STN disconnected from the pallidum in animal models would elucidate the role of these inter-structure connections and their necessity to ERNA. Furthermore, recordings of synaptic activity between the STN and GPe after persistent STN-DBS would indicate whether functional disconnection or vesicle depletion is present.

#### Vesicle Depletion is Necessary for Long-Term Dynamics of ERNA

Vesicle depletion was a fundamental addition to the model. Vesicle pool sizes can be estimated experimentally [12], and the evolution of the number of vesicles at synaptic terminals during cDBS could be compared to the vesicle depletion time-course predicted by our model. Alternatively, pharmacologically modulating vesicle re-uptake will modify the time constants of vesicle depletion. This in turn is expected to modulate the time constants of ERNA frequency decay, which could provide support for the role of vesicle depletion in the slow ERNA dynamics.

#### Key Features Required for Coupling Function

While methods to experimentally estimate PRCs are well established [55], estimating coupling functions between neurons is an area of active investigation [56]. As an alternative to estimating the coupling function, a micro electrode study of multiple STN neurons could be used to track firing times. Observing neuron firing times grouped into several clusters off stimulation would provide support for a coupling function with several stable zero-crossing points.

#### High Natural Frequency of Neurons is Necessary for the Frequency Decay

It is a fundamental property of the model that the oscillators have an intrinsic ability to fire at much higher rates than the off-stimulation network firing rate. We therefore predict that neurons that produce ERNA can fire at much higher frequencies in a fully disconnected state.

### Study Limitations

#### Fast Dynamics of Vesicle Depletion

The model in its current form is unable to replicate the characteristics of ERNA that are associated with the fast dynamics of vesicle depletion (see Supplementary Material sections C.1 and C.2). The model parameters were fitted to the characteristics of ERNA slow dynamics, which had previously eluded all modelling attempts. It is therefore not surprising that the model’s fast dynamics response does not line up as closely with the data as the long-term characteristics. In the data, we see growing inter-pulse evoked potential amplitude over the first 10 pulses before the amplitude steadies. Following termination of stimulation, the high amplitude evoked potentials behave similarly to a damped oscillation returning to its pre-stimulation levels over around 20ms [4]. We are able to capture some, but not all of the characteristics of the fast dynamics with our model. In particular, the inter-pulse evoked potentials do not decay between pulses and appear far more sinusoidal and noise-free than the data. Furthermore, oscillations following stimulation offset are not damped (see Supplementary Materials section C.1 for more details).

The model produces higher frequency evoked potentials with low-frequency stimulation, unlike the data which only showed high-amplitude, low-frequency responses to low-frequency stimulation [57, 58]. We demonstrate that if vesicle depletion did not occur with low-frequency stimulation, we would observe results similar to the data (see Supplementary Materials section C.2). We speculate that our current model does not replenish synaptic vesicles fast enough between pulses, which prevents us from capturing both the response to low-frequency stimulation, and the fast dynamics of ERNA as described above. In [6], there was little change in ERNA properties over time on adaptive deep brain stimulation (aDBS), unlike cDBS where ERNA properties change before reaching steady state. It was hypothesised that this could have represented different synaptic statuses due to the ability of synaptic pools to recover while on aDBS. Therefore, there may be more complicated synaptic dynamics in the early stages following stimulation onset. Modelling the biophysical details of the RRP dynamics, including the recycling of other vesicle pools into the RRP, may provide valuable versatility to the dynamics of the model and allow the model to reproduce these features.

#### Blind Validation of the Model

This study provides no fully blind validation of the modified Kuramoto model to the ERNA data. Despite reproducing a wide breadth of ERNA characteristics, the authors were not blinded to any of the data (even the characteristics used for validation, such as amplitude response in the variable neuromodulation sections). This is why this study is presented as a theoretical neuroscience investigation with predictions noted for future validation in other studies. However, we still manage to replicate a wide range of previously observed ERNA characteristics, a number of which could not be replicated by more complex previously published models.

#### Glutamatergic/GABAergic Properties of the Coupling Function

The fitted second-order coupling function is strongly negatively skewed (Fig 1E). This makes the biophysical interpretation of the coupling function more complex if considering a single, isolated population describing the STN, given that it is glutamatergic [59, 60]. However, even in the off-stimulation state the oscillators synchronise into clusters that receive phase-advancing effects from coupling, as the coupling function is slightly positive at a phase difference of zero (Fig 1E), indicating a glutamatergic tendency of the network. We interpret the negative shift of the coupling function as a phase indiscriminate GABAergic input from the GPe to the STN. Together, these characteristics manifests in the model as providing a phase advancing and glutamatergic (STN dominated) effect when oscillators are synchronised, but a phase delaying and GABAergic (GPe dominated) effect when the oscillators are not.

#### Other Biophysical Simplifications

We only simulate 50 coupled phase oscillators representing a larger neuronal network responsible for ERNA. While the number of phase oscillators is lowered for computational efficiency, the network’s behaviour approximates that of a larger neuronal population. Additionally, we do not consider the effect of different types of neurotransmitters. Instead we only model the impact of total vesicle availability on coupling strength. The interplay of multiple types of neurotransmitters and their dynamics between stimulation pulses may have to be considered to provide a more accurate representation of the oscillatory behaviour. Furthermore, the STN receives dynamic inputs from the GPe, cortex and other areas of the brain. Although we demonstrate here that only modelling one population’s dynamics is sufficient to replicate the slow dynamics of ERNA, including inputs from other neuronal structures may be key to reproducing the fast dynamics of ERNA.

## Conclusion

This work demonstrates that it is possible to replicate the long-term characteristics of ERNA by only considering the dynamics of a single neuronal structure. While the proposed computational model remains to be validated using follow-up data, this work has the potential to provide insights into the underlying mechanisms of ERNA. Biophysiological properties of the model key to reproduce ERNA are revealed to be vesicle depletion, properties of the coupling function, and high natural frequency of individual oscillators. These properties may inform future investigations into the underlying mechanisms of ERNA.

## Acknowledgements

JS and TD are supported by DARPA HR0011- 20-2-0028 Manipulating and Optimising Brain Rhythms for Enhancement of Sleep (Morpheus) and the UK Medical Research Council grant MC UU 00003/3. CW and HT are supported by the Medical Research Council (MC UU 00003/2, MR/V00655X/1, MR/P012272/1), the National Institute for Health Research (NIHR) Oxford Biomedical Research Centre (BRC), and the Rosetrees Trust, UK. BD is jointly supported by the Royal Academy of Engineering and Rosetrees under the Research Fellowship programme, and was also supported by Medical Research Council grant MC UU 00003/1. Content represents views of the authors and not the funders.

## Authorship Contributions

**JS** - Conceptualization, Formal analysis, Investigation, Methodology, Software, Validation, Visualization, Roles/Writing - original draft, Writing - Review and Editing **CW** - Methodology, Writing – Review & Editing **HT** - Funding Acquisition, Supervision, Writing – Review and Editing **TD** - Funding Acquisition, Supervision, Writing – Review and Editing **BD** - Conceptualization, Formal Analysis, Funding, Acquisition, Investigation, Methodology, Software, Supervision, Validation, Visualization, Writing – Review and Editing

## Supplementary Material

### A Fitted Model Parameter Set

The results of the fitting process (outlined in Methodology section *Fitting Process*) are presented in Table A. All model parameters presented in the top row of Table A were optimisable parameters for the fitting procedure. Some of the parameters were additionally subject to manual parameter tuning in order to provide model dynamics that resembled the data more closely. However, this represented only small adjustments by comparison to their fitted value. The bottom row of parameters in Table A are parameters of the three vesicle pool model and were largely determined from data as detailed below, with a small amount of manual parameter tuning within the bounds set by the data.

**Table A.**
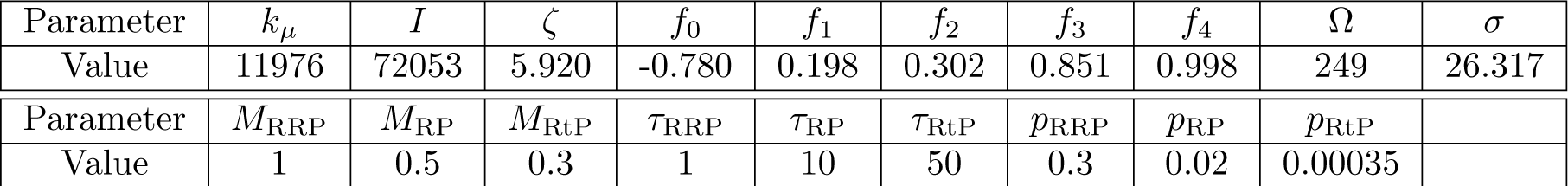
Fitted model parameter set. The base model parameter set for all simulations. The bottom row are parameters of the three vesicle pool model.

| Parameter | $k_\mu$ | $I$ | $\zeta$ | $f_0$ | $f_1$ | $f_2$ | $f_3$ | $f_4$ | $\Omega$ | $\sigma$ |
| --- | --- | --- | --- | --- | --- | --- | --- | --- | --- | --- |
| Value | 11976 | 72053 | 5.920 | -0.780 | 0.198 | 0.302 | 0.851 | 0.998 | 249 | 26.317 |
| Parameter | $M_{\text{RRP}}$ | $M_{\text{RP}}$ | $M_{\text{RtP}}$ | $\tau_{\text{RRP}}$ | $\tau_{\text{RP}}$ | $\tau_{\text{RtP}}$ | $p_{\text{RRP}}$ | $p_{\text{RP}}$ | $p_{\text{RtP}}$ | |
| Value | 1 | 0.5 | 0.3 | 1 | 10 | 50 | 0.3 | 0.02 | 0.00035 |  |

The time constant of each pool *τ*_pool_ is estimated using the approximate time required to replenish the vesicle pool through recycling in the absence of unphysiological stimulation. The RRP recycles quickly, on the order of seconds [12], we therefore chose *τ*_RRP_ to be 1 second. The RP also recycles quickly on the order of seconds [12], and given the size difference between the RRP and the RP, *τ*_RRP_ is taken to be 10 seconds. The RtP takes longer to replenish recycling on the order of minutes [12], *τ*_RRP_ is therefore chosen to be 50 seconds.

The probability of vesicle release by a stimulation pulse *p*_pool_ is estimated from the time taken to deplete each vesicle. The RRP is expected to deplete within 10 pulses of unphysiological stimulation [13], the RP over seconds and the RtP over minutes [12]. The value of *p*_pool_ is taken to be inversely proportional to the number of pulses required to deplete each pool. The proportionality constant is set in combination with *τ*_pool_ according the stable occupancy of each pool on 130Hz stimulation. Therefore, *p*_RRP_ is given a value of 0.3 (corresponding to 9 pulses on 130Hz stimulation), *p*_RP_ a value of 0.02 (corresponding to one second on 130Hz stimulation or 130 pulses) and *p*_RtP_ a value of 0.0035 (one minute or 7800 pulses) to align with these observations.

The estimated contribution of each vesicle pool on coupling strength, *M*_pool_, is modelled as proportional to the product of the relative size of each vesicle pool (*N*_pool_) and the inverse of the vesicle pools distance from the synaptic terminal (*d*_pool_),

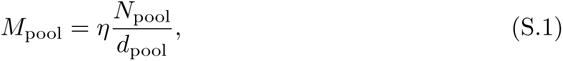

where *η* is a normalising factor, the same across vesicle pools.

The RRP is the smallest vesicle pool, usually approximated to be between 1 − 2% of the overall number of vesicles [12]. Additionally, RRP vesicles are usually considered to be either docked or very close to the synaptic terminal. Given the approximate size of vesicle pool around 20nm [39, 40] in diameter, we estimate the average distance of RRP vesicles to the synaptic terminal to be on the scale of 10nm. The important consideration for *N*_pool_ is the relative size between pools. We therefore take the total number of vesicles as 100, and taking the percentage of vesicles in the RRP as 1%, we obtain *N*_RRP_*/d*_RRP_ = 0.1nm*^−^*^1^. We choose the normalising factor *v* = 10nm (across all pools), which gives *M*_RRP_ = 1. The RP contains slightly more vesicles at around 10 − 20% [12], but is located further from the synaptic terminal. Furthermore, the RP must release fewer vesicles to the synapse when full compared to the RRP. Electron microscopy and neurotransmitter release simulations suggests that the majority of vesicles located proximal to the synapse are located within 500nm of the terminal [12, 36, 37, 40], therefore we estimate the average distance of the RP vesicles to the synaptic terminal as 200nm. Taking the percentage of vesicles in the RP as 10%, we obtain *N*_RP_*/d*_RP_ = 0.05nm*^−^*^1^ and *M*_RP_ = 0.5. Lastly, the RtP represents the majority of vesicles, 80 − 90% [12], and given the percentages taken for the other vesicle pools we will assume the RtP represents 89%. Estimates for the furthest distance vesicle pools take from the synaptic terminal are between 1_µ_m [40] and 5_µ_m [12, 38], we therefore take an approximate distance of 3µm. We obtain *M*_RtP_ = 0.3.

### B Explaining Model Dynamics

#### B.1 Amplitude Response

One of the benefits of using computational models is that we have the ability to investigate the dynamics that may be responsible for the ERNA characteristics in the simulations. Here we investigate what causes the amplitude variation seen over the longer time period. By recording the phase difference between one random oscillator against all the others, we are able to observe clustering behaviour over both the off- and on-stimulation periods (Fig S.1). Fig S.1Bi shows that in the steady off-stimulation state we see two clusters approximately anti-phase (due to the coupling function, see Fig 1E), which result in low amplitude activity. In on-stimulation histograms of Figs S.1Bii, iii and iv, it can be seen that there is a shift away from an even distribution between each cluster, with a preference to synchronise to one cluster. The development of this distribution of phases between clusters over the stimulation window accounts for the amplitude variation. Firstly, the histograms of Figs S.1Biii and iv do not differ greatly as can be seen from the amplitude dynamics with little variation after approximately 50 seconds of stimulation. In the early part of the stimulation window there is greater coupling around a single cluster at an approximate ratio of 25:1 oscillators in each group compared to the smaller cluster. In the later part of the stimulation window, this ratio shrinks to around 10:1 as coupling strength drops further and there is less attraction to stay coupled to the dominant cluster.

#### B.2 Frequency Response

The on-stimulation frequency response can be explained by investigating the dynamics of individual oscillators as well. After each stimulation pulse, the oscillators are pushed towards synchronisation around a single phase due to the sinusoidal PRC. This would occur for any PRC with a stable zero crossing point. Furthermore, this occurs regardless of the length of time following stimulation onset as this is a property that is independent of vesicle depletion’s effect on coupling strength. Early in the stimulation window, before vesicle depletion has reduced coupling strength too greatly, the oscillators remain mostly synchronised around a single phase for the majority of the remaining inter-pulse interval (see Fig S.1Bii). Given that coupling strength is still high and that the coupling function is positive for zero phase difference, this increases the frequency of the network at this early stage. Later in the stimulation window, as coupling strength reduces, fewer oscillators remain synchronised and for a shorter period of time (see Fig S.1Biii and iv). As coupling strength is much weaker, there is less of a contribution to oscillator phase advancement from the interaction with other oscillators.

**Fig S.1.**
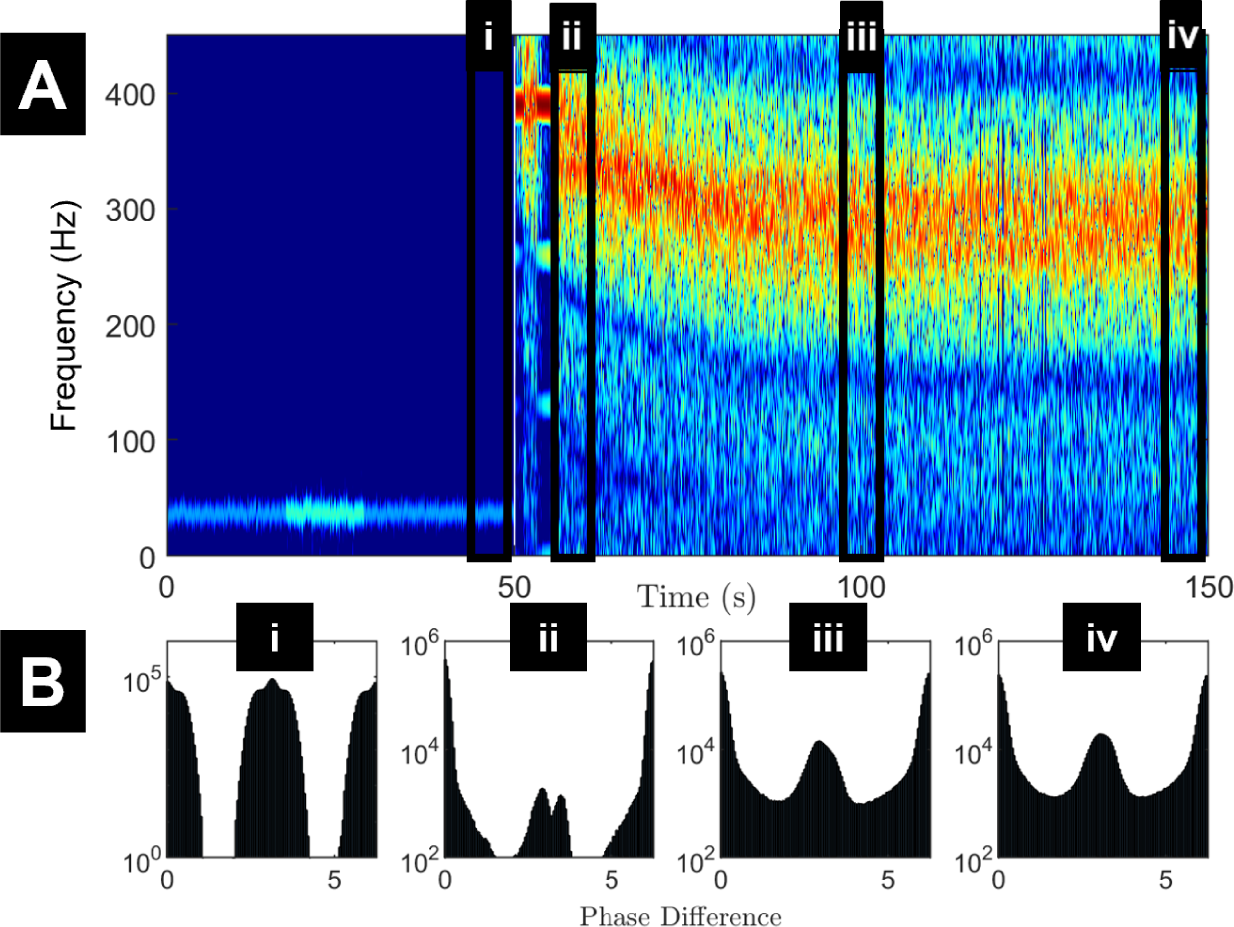
Clustering of oscillators explains the amplitude dynamics over continuous stimulation. (A) Spectrogram of network activity (Same as in Fig 1A) over 50 seconds off-stimulation before turning on-stimulation for 100 seconds. The black boxes labelled i, ii, iii and iv refer to the periods clustering was observed over for the histograms in panel B. (B) Histograms of phase difference for four five second periods. (i) 44 to 49 seconds, (ii) 55 to 60 seconds (to avoid brief period of entrainment), (iii) 97.5 to 102.5 seconds, and (iv) 144 to 149 seconds. (i) Plotted with a y axis limit of 10^0^ to 10^6^, whereas the other a plotted with a limit of 10^2^ to 10^6^, to demonstrate the lack of activity observed outside the clusters.

As the stimulation PRC integrates to zero and noise is drawn from a random distribution with a mean of zero, both of these potential phase effecting mechanism trend to having zero net effect with enough time. Hence, with a much lower coupling strength, the primary effect on oscillator phase progression comes from the natural frequency of the individual oscillators (Eq.1). Therefore the network trends to a frequency close to the fitted mean of the natural frequencies of all the oscillators in the network after a long period of high frequency stimulation (see Ω = 249 in Table A).

### C Current Model Shortcomings

#### C.1 Fast Dynamics

For comparison to the fast ERNA dynamics reported in [4], DBS is turned on at 130Hz for a period of 100ms with 50ms of observations off-stimulation either side of the DBS window. This follows an initial 10 seconds of the model settling. Initially coupling strength decayed on-stimulation following our three vesicle pool model with the parameters obtained from the fitting process (Table A). Coupling strength begins to recover with a one second time constant as dictated by RRP recovery. However, high frequency and high amplitude dynamics are maintained during the 100ms following stimulation termination due to coupling strength remaining low at this time scale (Fig S.2). Model dynamics return to low frequency conditions within around one second following stimulation termination (see for example Fig 2B). We also note that inter-pulse dynamics are very regular and do not resemble typical ERNA signals. Hence, our fitted model reproduces the slow dynamics of ERNA, but is unable to capture its fast dynamics. This may be due to the slow replenishment of vesicles in our simplified model on this time scale, and introducing more complex dynamics such as vesicle mixing between pools or STN-GPe excitatory/inhibitory connections may be necessary to reproduce fast ERNA dynamics.

**Fig S.2.**
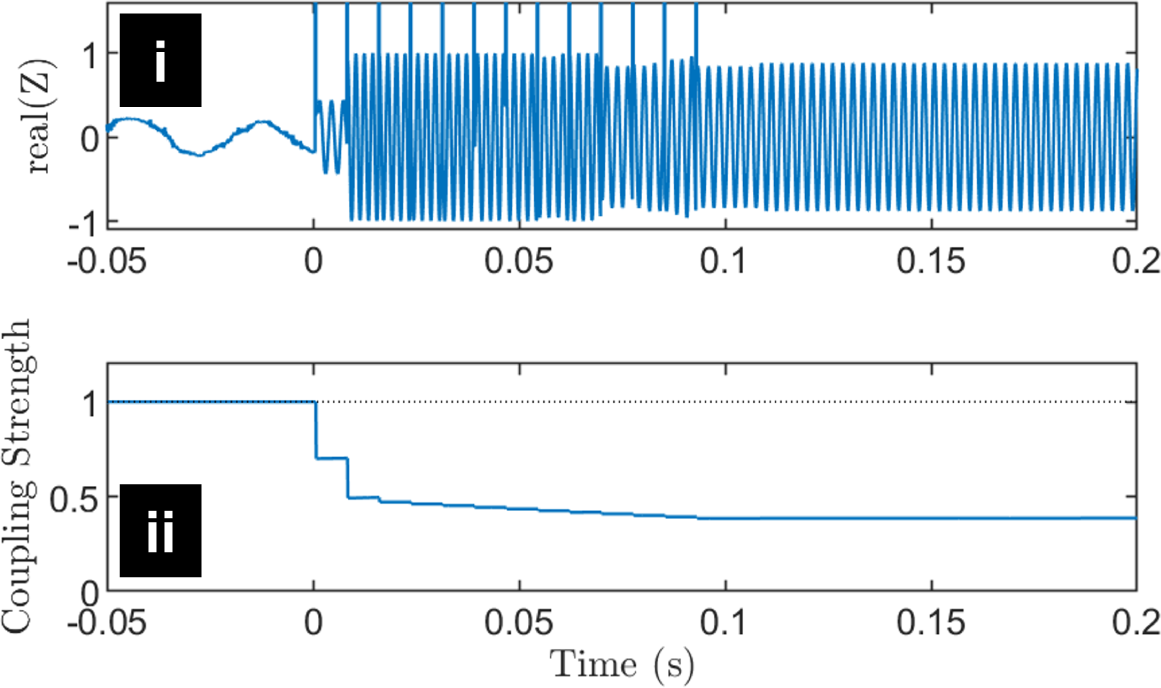
Fast dynamics of ERNA for 100ms of 130Hz stimulation pulses. (i) The approximate amplitude response with stimulation artefact artificially superimposed on the signal following the simulation for demonstration purposes. (ii) The corresponding coupling strength of the network.

#### C.2 Subtherapeutic Stimulation

Subtherapeutic stimulation at 20Hz is capable of eliciting evoked potentials [57]. Similar responses were also seen at lower stimulation frequencies of 3Hz [58]. These evoked potentials had increased amplitude compared to baseline, but no high frequency content. 20Hz stimulation may be considered on the borderline of physiological-unphysiological stimulation [14], while 3Hz stimulation is certainly within the physiological limits. However, this is dependent on the neural structure being stimulated and 20Hz may even be considered within physiological frequencies in some structures, especially if this stimulation frequency is close to the natural frequency of the neuronal ensemble.

The application of low frequency stimulation to our model does not align with these results, however. For both 3Hz and 20Hz we still see the promotion of high frequency activity (Figs S.3A and E). Stimulation at 3Hz produces evoked responses at lower frequencies compared to higher stimulation frequencies tried. Hints from literature can help us investigate this shortcoming further. Stimulation at 10Hz has been shown to not cause vesicle depletion in some cases [11]. Hence, it is likely that little vesicle depletion will be observed in response to 20Hz stimulation. Furthermore, as 20Hz stimulation elicits very similar responses to 3Hz stimulation [58], it could be expected that these two stimulation paradigms elicit similar neural responses. Hence, we investigate the evoked response in the absence of vesicle depletion for 20Hz stimulation (Fig S.3I), otherwise the model and parameters remain unchanged.

Without vesicle depletion in the model, we do not see the evolution of high frequency activity (see Figs S.3I and L). However, we still see the promotion of high amplitude activity, which is in line with [57, 58] (see Fig S.3K). This is primarily due to the maintenance of a higher coupling strength from the lack of vesicle depletion. Therefore, it is likely that improving the fast dynamics of the model as suggested in the previous section (especially with regards to the dynamics of the RRP) would make the model behave more similarly to data in response to subtherapeutic stimulation.

### D Fitting Robustness

Here we explore how changing the coupling function and the PRC in the fitted model affects the ERNA response to stimulation.

#### D.1 Sensitivity of Coupling Function

A second-order Fourier series coupling function was able to capture more complex neuronal interactions than the sine function commonly used in the Kuramoto model. As shown in the main text, fitting the Fourier series coefficients provided slow dynamics closely approximating the ERNA spectrogram. For comparison we also simulate the model with a sinusoidal coupling function and a coupling function derived from the Hodgkin-Huxley neuron model [55], while all other parameters remain constant (see Fig S.4). These coupling functions do not lead to ERNA dynamics (Figs S.4B and C). Additionally, the dynamics of the sine coupling function result in activity at a much higher frequency off stimulation (around 200-250Hz), and at a much higher amplitude (Fig S.4B). This is due to simpler clustering behavior compared to the fitted coupling function, which favors two clusters in the absence of stimulation. The sine coupling function does not give rise to changes in oscillation frequency with stimulation despite vesicle depletion remaining present (Figs S.4B). The Hodgkin-Huxley derived coupling function shows a response where frequency increases with time on-stimulation and vesicle depletion (Figs S.4C). While these results are based on model parameters obtained for the second-order Fourier coupling function, attempts to fit to the long-term ERNA frequency decay using each of these coupling functions were made, but no promising results were obtained.

The Fourier series fitted coupling function is a key element of the model to reproduce the ERNA frequency decay, therefore we also investigate the sensitivity of the five fitted coefficient values. For each of the Fourier series coefficients we re-simulate the computational model after modulating the coefficients by the factors 0, 0.25, 0.5, 1, 2 and 4 (Fig S.5). The columns corresponding to the first-order coefficients (*f*_1_ and *f*_2_) show little variability for all factors (including 0) both on and off-stimulation. Hence, the first order coefficients are not only not very sensitive, but may not be necessary for the model. The other three coefficients all seem to produce more notable changes to the spectrograms when modulated. The changes result in either failing to reproduce off-stimulation dynamics, on-stimulation dynamics, or both such that the resulting spectrogram no longer matches all characteristics of the slow ERNA dynamics.

#### D.2 Sensitivity of Phase Response Curve

We also compare different types of PRCs (Fig S.6). We scale the Hodgkin-Huxley PRC to a maximum amplitude of one to provide a comparable scale to the sinusoidal PRC. We consider a constant PRC of magnitude 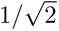. These PRCs are chosen so that we can identify the role of the PRC in the computational model. As the constant PRC does not replicate the on-stimulation dynamics, it is likely that a synchronising effect from the PRC is required to reproduce ERNA. This is in agreement with the results from Supplementary Materials section *Explaining the Amplitude Response*. The Hodgkin-Huxley PRC (Fig S.6B) is chosen as a more biophysical representation, and we do see the peak of the frequency decay start at 390Hz (the third harmonic of stimulation frequency) before dropping down. There is intermittent locking at the 5:2 harmonic (325Hz for 130Hz stimulation frequency) before dropping further and locking to the second harmonic of stimulation (at 260Hz). However, unlike the sine PRC and the data, the locking to components of stimulation frequency is much stronger to the point where activity only seems to appear at these components. This could be due to the sharper gradient of the Hodgkin-Huxley PRC at the zero crossing point having a stronger synchronising effect on the oscillators than the sinusoidal PRC. As the fitting was performed using the sinusoidal PRC, it is certainly possible that by fitting with the Hodgkin-Huxley PRC instead we could observe a response that more closely resembles the data.

### E Alternative Parameter Set with Lower Natural Frequency

We explore the modified Kuramoto model further by constraining the natural frequencies of individual oscillators to lower frequencies. Results from STN slice experiments could indicate that individual STN oscillators have natural frequencies around 20Hz. Previously, we assumed that 20Hz was the frequency of the overall neural ensemble and didn’t reflect individual neuron properties, therefore not constraining the *ω_i_*’s in the model. Here, we constrain the mean natural frequency of oscillators (Ω) to 20Hz. Given the model set up, it is possible to maintain similar off-stimulation dynamics by countering the decrease in natural frequency by a corresponding increase in the noughth order shift of the coupling function *f*_0_ (see values in Table B).

**Table B.** Alternative parameter sets for low natural frequency simulations. All other parameters remain unchanged from Table A.

| Parameter | $\Omega$ | $\sigma$ | $f_0$ | $p_{\text{RtP}}$ | $\tau_{\text{RtP}}$ |
| --- | --- | --- | --- | --- | --- |
| Values for Fig S.7 | 20 | 2.11 | -0.68 | 0.00035 | 50 |
| Values for Fig S.8 | 20 | 2.11 | -0.68 | 0.00007 | 10 |

Following the adjustments to natural frequency and *f*_0_, we simulate the model while keeping all other parameters the same as the fitted model (Fig S.7). We see similar amplitude dynamics decreasing over the stimulation window (Fig S.7C) as with the fitted model. However, as coupling strength continues to decrease over the stimulation window we see the frequency decay reach much lower values towards the end of the stimulation window (Fig S.7B). It continues to decrease to a value lower than that seen from the ERNA data (*<*100Hz).

Given how vesicle depletion changes the coupling strength during stimulation, it is not possible to maintain the same dynamics on-stimulation with this same method. Hence, in order to maintain a similar frequency decay, the long term dynamics of the RtP (as dictated by *τ*_RtP_ and *p*_RtP_) are modulated (see second row of Table B) such that the frequency at the end of the 100-second stimulation window approximately matches that of the fitted modified Kuramoto model from the main part of this study (Fig 1)A. While it is possible to maintain higher frequency dynamics (Fig S.8B), the amplitude decay does not demonstrate any substantial variation, remaining somewhat constant (Fig S.8C). Hence, in order to see amplitude variation it may be required that RtP occupancy and therefore coupling strength decreases low enough such that oscillators can move between clusters. By reducing coupling strength closer to zero we allow the network to oscillate at a similar frequency to the natural frequency of the individual oscillators. While it stands to reason that this could be achieved with an alternative resolution of the coupling function, we have explored this hypothesis in detail. We have repeated the fitting process with constrained natural frequencies to 20Hz and have not observed encouraging results.

**Fig S.3.**
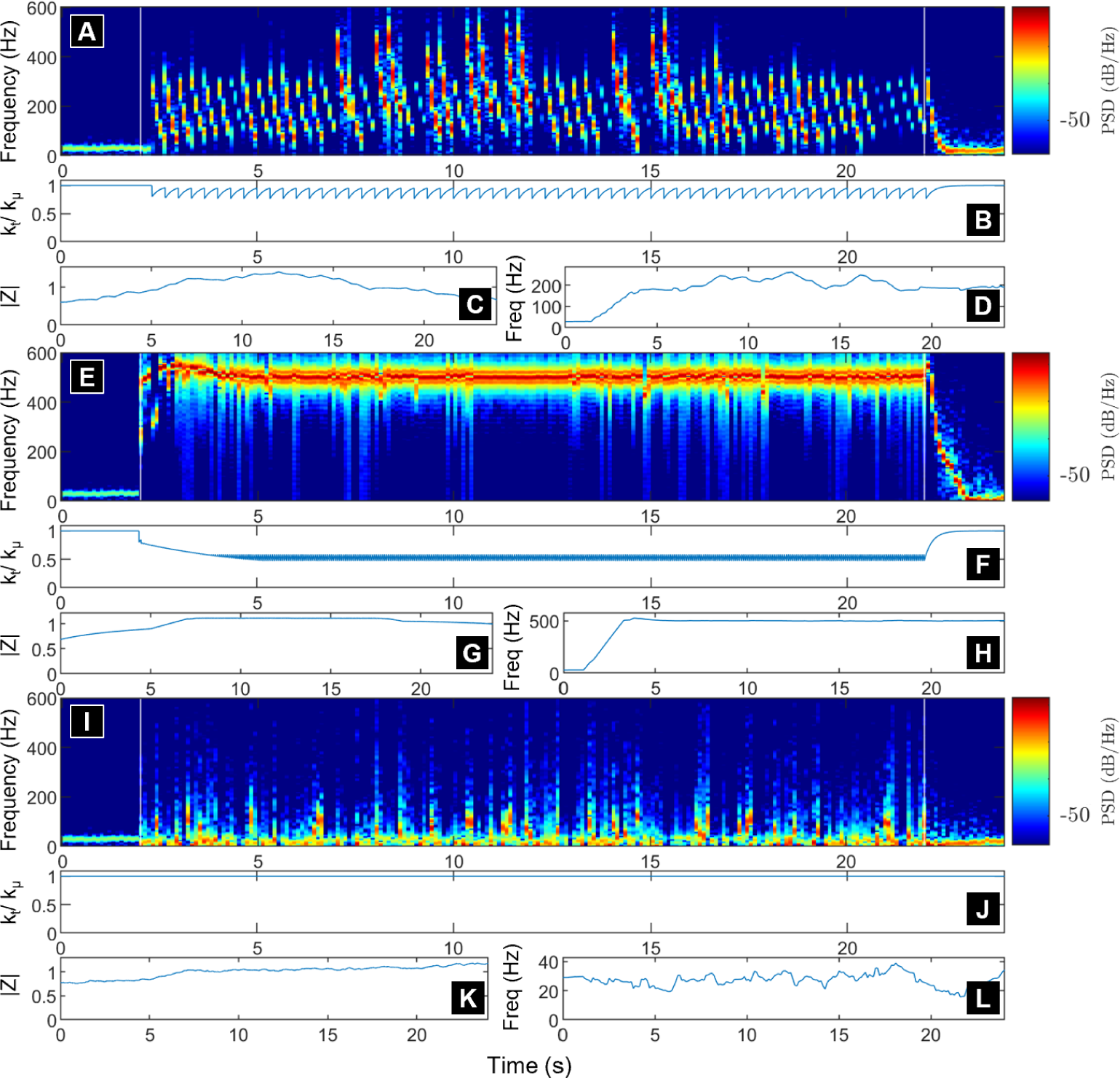
Response of the model to low frequency stimulation with and without vesicle depletion as a theoretical mechanism of subtherapeutic stimulation. (A, E and I) Spectrogram with stimulation frequency at 3Hz (A) and 20Hz (E and I). Stimulation onset at two seconds into simulation indicated by white line. (B, F and J) The evolution of coupling strength for each simulation. (C, G and K) Amplitude of oscillations over whole simulation. (D, H and L) Peak frequency response for each of the 100ms windows over the whole simulation. (A-H) Vesicle depletion consistent with Eqs. 4 and 5. (I-L) No vesicle depletion.

**Fig S.4.**
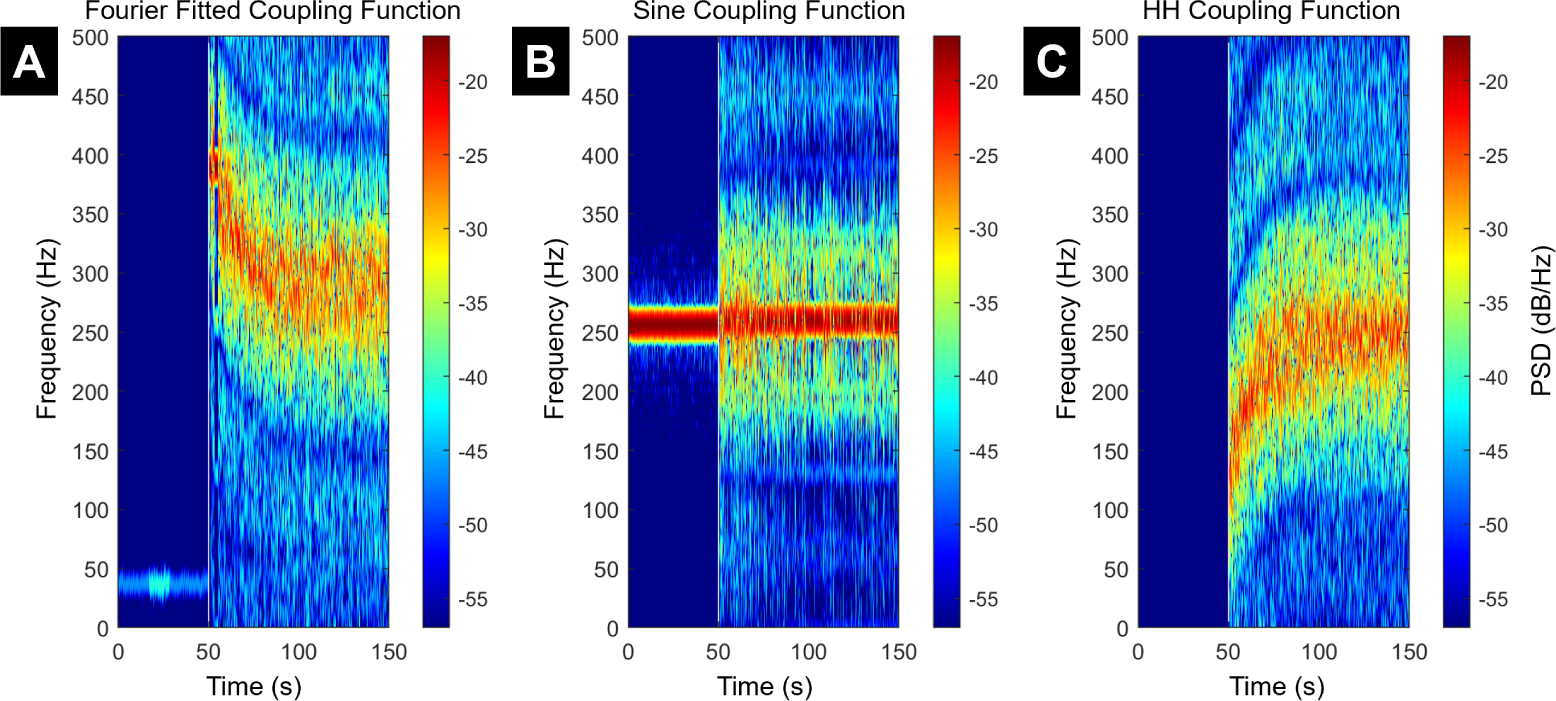
Effect of the coupling function on ERNA slow dynamics. 50 seconds off-stimulation followed by 200 seconds on-stimulation. (A) The fitted second-order Fourier series coupling function used in the rest of the study, similar figure to Fig 1A. (B) Sinusoidal coupling function. (C) Hodgkin-Huxley (HH) derived coupling function. Other than coupling function changes, all other model parameters remain the same.

**Fig S.5.**
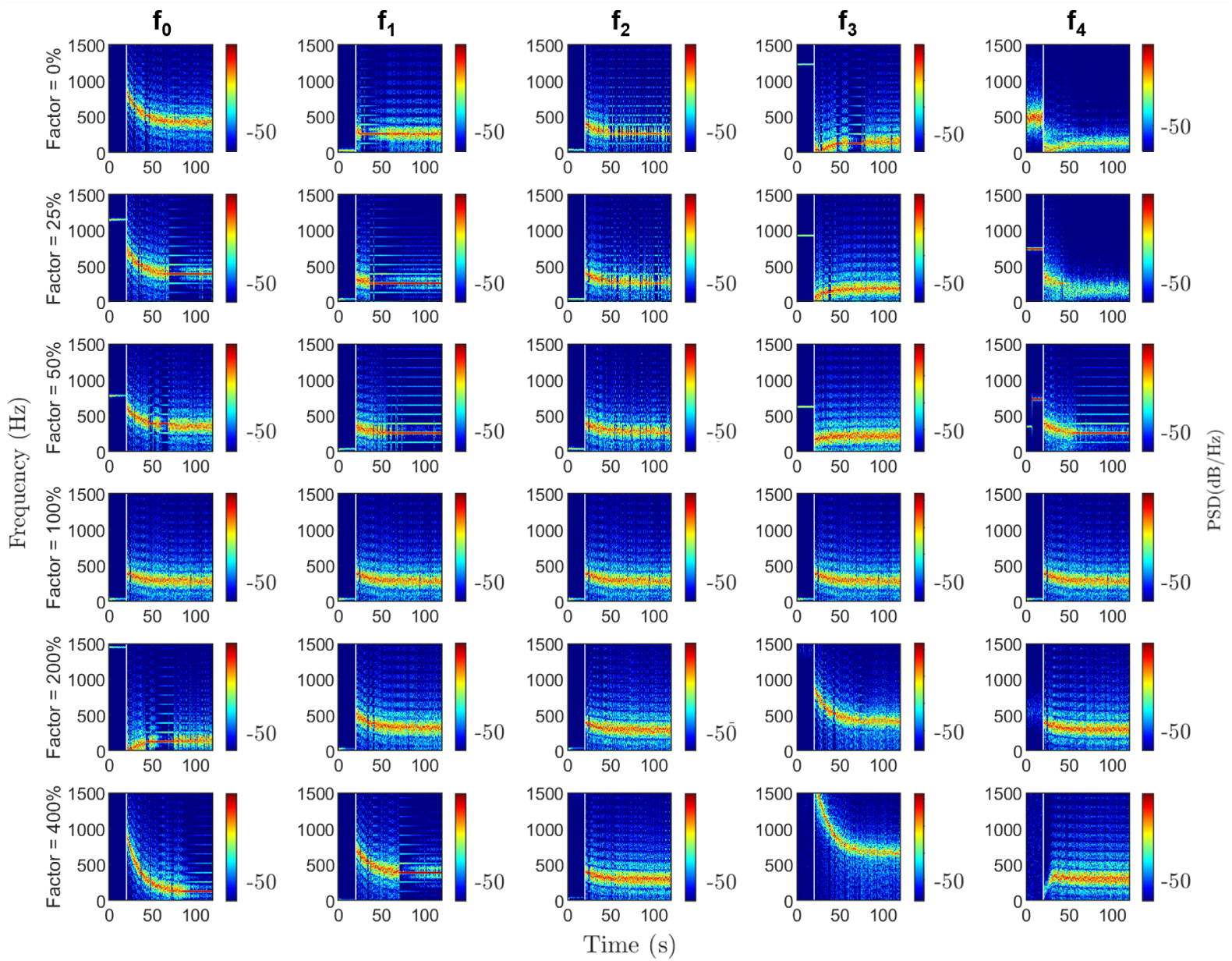
Effect of modulating each Fourier coefficient of the coupling function on ERNA slow dynamics. Spectrograms showing 20 seconds off-stimulation followed by 100 seconds on-stimulation, observed over the large frequency range of [0Hz,1500Hz] due to high frequency activity in some of the simulations. Columns ordered by Fourier coefficients: *f*_0_ correspond to the vertical shift, *f*_1_ and *f*_2_ are coefficients corresponding to the first harmonic components (cosine and sine, respectively), and *f*_3_ and *f*_4_ are coefficients corresponding to the second harmonic components (cosine and sine, respectively). Rows ordered by the factor multiplying the coefficients. Factor of 100% represents no change to the coefficient with respect to the fitted results, so this row has the same parameters for each of the coefficient simulations.

**Fig S.6.**
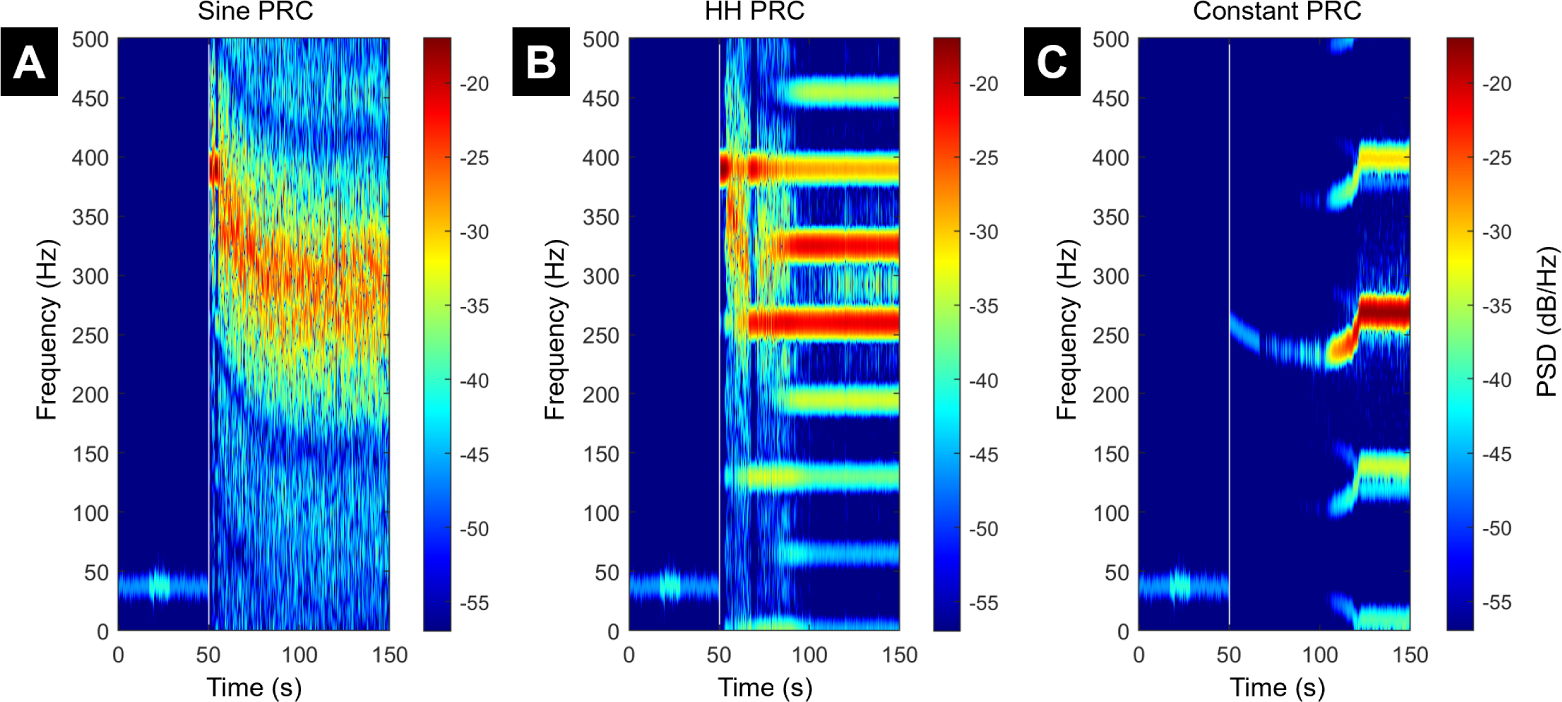
Effect of the PRC on ERNA slow dynamics. 50 seconds off-stimulation followed by 200 seconds on-stimulation. (A) sinusoidal PRC, similar simulation to Fig 1A. (B) Hodgkin-Huxley (HH) PRC. (C) Constant PRC. Other than PRC changes all parameters remain the same as the main body of this study.

**Fig S.7.**
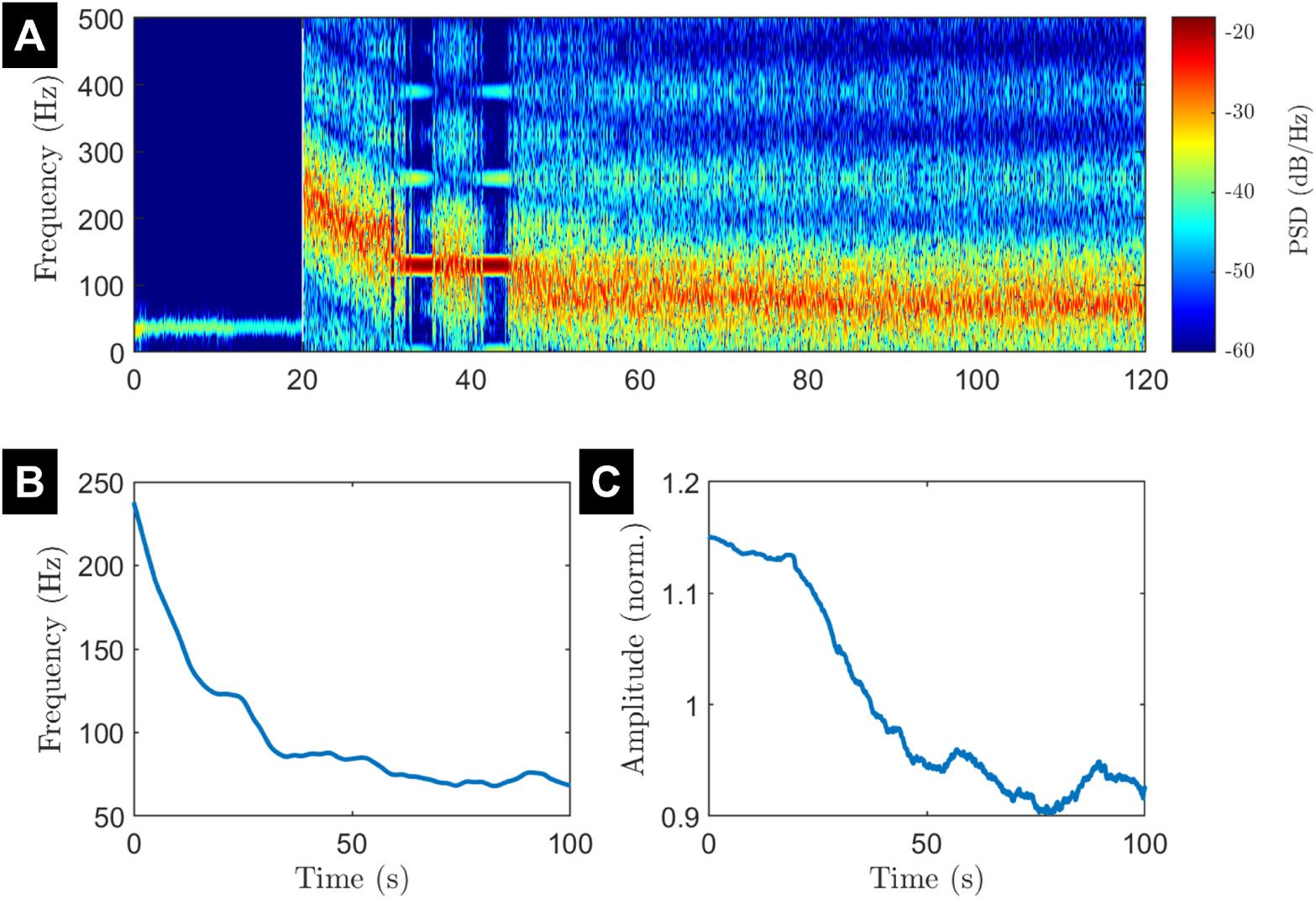
Slow dynamics for lowered natural frequency model with all other parameters remaining the same. (A) The spectrogram of 20 seconds off-stimulation followed by 100 seconds on-stimulation. (B) The on-stimulation frequency decay. (C) the on-stimulation amplitude response.

**Fig S.8.**
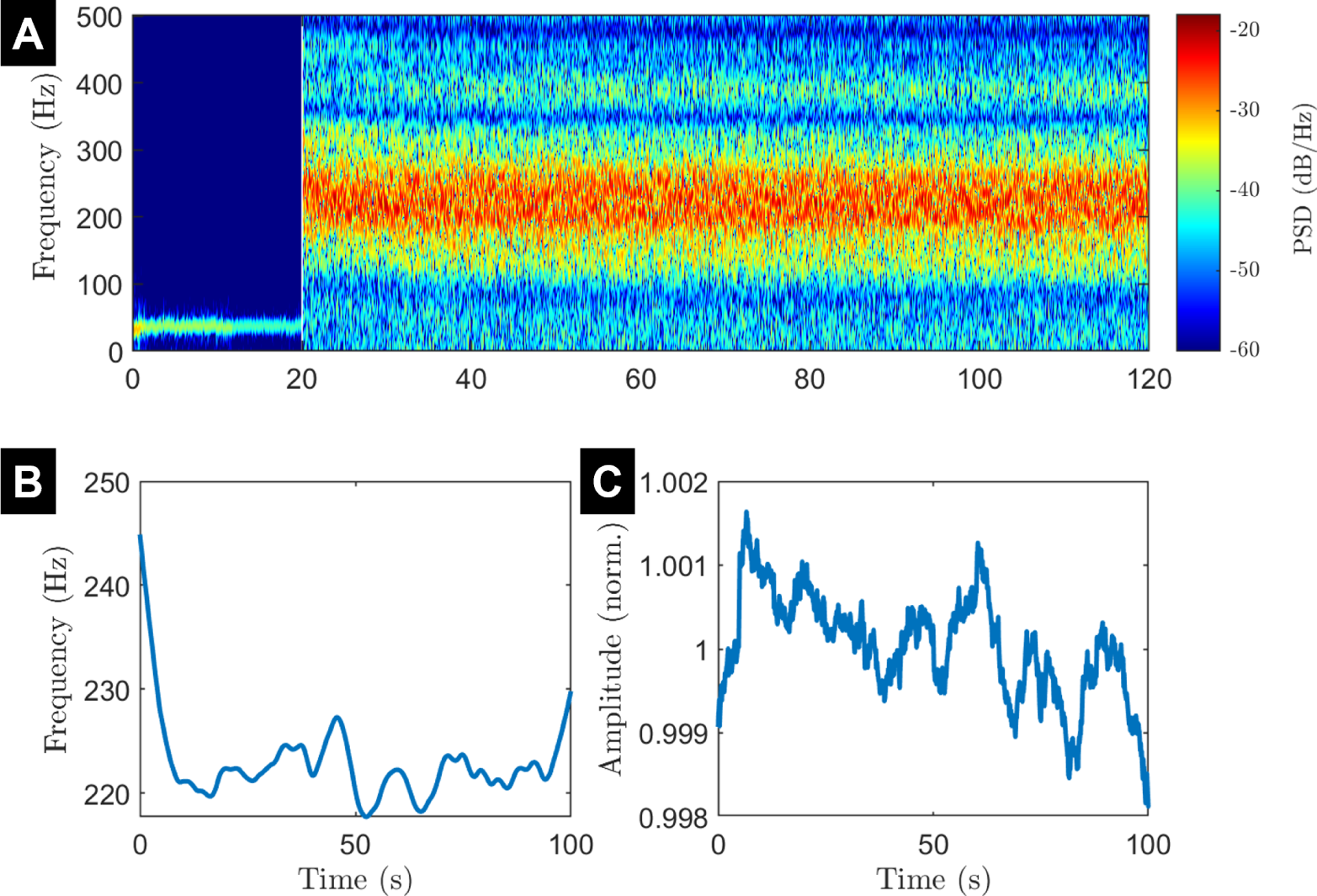
Slow dynamics for lowered natural frequency model with coupling strength changed to approximately fit to frequency decay. (A) The spectrogram of 20 seconds off-stimulation followed by 100 seconds on-stimulation. (B) The on-stimulation frequency decay. (C) the on-stimulation amplitude response.

## References

1. Wiest C, Tinkhauser G, Pogosyan A, Bange M, Muthuraman M, Groppa S, et al. Local field potential activity dynamics in response to deep brain stimulation of the subthalamic nucleus in Parkinson’s disease. Neurobiology of Disease. 2020;143. doi:10.1016/j.nbd.2020.105019.

2. Schmidt SL, Brocker DT, Swan BD, Turner DA, Grill WM. Evoked potentials reveal neural circuits engaged by human deep brain stimulation. Brain Stimulation. 2020;13. doi:10.1016/j.brs.2020.09.028.

3. Johnson KA, Cagle JN, Lopes JL, Wong JK, Okun MS, Gunduz A, et al. Globus pallidus internus deep brain stimulation evokes resonant neural activity in Parkinson’s disease. Brain Communications. 2023;5. doi:10.1093/braincomms/fcad025.

4. Sinclair NC, McDermott HJ, Bulluss KJ, Fallon JB, Perera T, Xu SS, et al. Subthalamic nucleus deep brain stimulation evokes resonant neural activity. Annals of Neurology. 2018;83:1027–1031. doi:10.1002/ana.25234.

5. Sinclair NC, Fallon JB, Bulluss KJ, Thevathasan W, McDermott HJ. On the neural basis of deep brain stimulation evoked resonant activity. Biomedical Physics and Engineering Express. 2019;5. doi:10.1088/2057-1976/ab366e.

6. Wiest C, He S, Duchet B, Pogosyan A, Benjaber M, Denison T, et al. Evoked resonant neural activity in subthalamic local field potentials reflects basal ganglia network dynamics. Neurobiology of Disease. 2023;178. doi:10.1016/j.nbd.2023.106019.

7. Chiken S, Nambu A. Mechanism of Deep Brain Stimulation: Inhibition, Excitation, or Disruption? Neuroscientist. 2016;22. doi:10.1177/1073858415581986.

8. Milosevic L, Kalia SK, Hodaie M, Lozano AM, Fasano A, Popovic MR, et al. Neuronal inhibition and synaptic plasticity of basal ganglia neurons in Parkinson’s disease. Brain. 2018;141. doi:10.1093/brain/awx296.

9. Steiner LA, Tomás FJB, Planert H, Alle H, Vida I, Geiger JRP. Connectivity and dynamics underlying synaptic control of the subthalamic nucleus. Journal of Neuroscience. 2019;39. doi:10.1523/JNEUROSCI.1642-18.2019.

10. Xu SS, Lee WL, Perera T, Sinclair NC, Bulluss KJ, McDermott HJ, et al. Can brain signals and anatomy refine contact choice for deep brain stimulation in Parkinson’s disease? Journal of Neurology, Neurosurgery and Psychiatry. 2022;93. doi:10.1136/jnnp-2021-327708.

11. Fernández-Alfonso T, Ryan TA. The kinetics of synaptic vesicle pool depletion at CNS synaptic terminals. Neuron. 2004;41. doi:10.1016/S0896-6273(04)00113-8.

12. Rizzoli SO, Betz WJ. Synaptic vesicle pools. Nature Reviews Neuroscience. 2005;6. doi:10.1038/nrn1583.

13. Kaeser PS, Regehr WG. The readily releasable pool of synaptic vesicles. Current Opinion in Neurobiology. 2017;43. doi:10.1016/j.conb.2016.12.012.

14. Denker A, Rizzoli SO. Synaptic vesicle pools: An update. Frontiers in Synaptic Neuroscience. 2010;doi:10.3389/fnsyn.2010.00135.

15. Alabi ARA, Tsien RW. Synaptic vesicle pools and dynamics. Cold Spring Harbor Perspectives in Biology. 2012;4. doi:10.1101/cshperspect.a013680.

16. Hallworth NE, Wilson CJ, Bevan MD. Apamin-sensitive small conductance calcium-activated potassium channels, through their selective coupling to voltage-gated calcium channels, are critical determinants of the precision, pace, and pattern of action potential generation in rat subthalamic nucleus neurons in vitro. Journal of Neuroscience. 2003;23. doi:10.1523/jneurosci.23-20-07525.2003.

17. Wilson CJ, Weyrick A, Terman D, Hallworth NE, Bevan MD. A Model of Reverse Spike Frequency Adaptation and Repetitive Firing of Subthalamic Nucleus Neurons. Journal of Neurophysiology. 2004;91. doi:10.1152/jn.00924.2003.

18. Kitai ST, Kita H. Anatomy and Physiology of the Subthalamic Nucleus: A Driving Force of the Basal Ganglia. In: Carpenter MB, Jayaraman A, editors. The Basal Ganglia II. Boston, MA: Springer US; 1987. p. 357–373.

19. Schor JS, Montalvo IG, Spratt PWE, Brakaj RJ, Stansil JA, Twedell EL, et al. Therapeutic Deep Brain Stimulation Disrupts Movement-Related Subthalamic Nucleus Activity in Parkinsonian Mice. eLife. 2022;11. doi:10.7554/elife.75253.

20. Tass PA, Majtanik M. Long-term anti-kindling effects of desynchronizing brain stimulation: A theoretical study. Biological Cybernetics. 2006;94(1):58–66. doi:10.1007/s00422-005-0028-6.

21. Asllani M, Expert P, Carletti T. A minimally invasive neurostimulation method for controlling abnormal synchronisation in the neuronal activity. PLoS Computational Biology. 2018;14(7):e1006296. doi:10.1371/journal.pcbi.1006296.

22. Weerasinghe G, Duchet B, Cagnan H, Brown P, Bick C, Bogacz R. Predicting the effects of deep brain stimulation using a reduced coupled oscillator model. PLoS Computational Biology. 2019;15. doi:10.1371/journal.pcbi.1006575.

23. Bick C, Goodfellow M, Laing CR, Martens EA. Understanding the dynamics of biological and neural oscillator networks through exact mean-field reductions: a review. The Journal of Mathematical Neuroscience. 2020;10(1):9. doi:10.1186/s13408-020-00086-9.

24. Weerasinghe G, Duchet B, Bick C, Bogacz R. Optimal closed-loop deep brain stimulation using multiple independently controlled contacts. PLOS Computational Biology. 2021;17(8):e1009281. doi:10.1371/journal.pcbi.1009281.

25. Duchet B, Sermon JJ, Weerasinghe G, Denison T, Bogacz R. How to entrain a selected neuronal rhythm but not others: open-loop dithered brain stimulation for selective entrainment. Journal of Neural Engineering. 2023;20(2):026003. doi:10.1088/1741-2552/acbc4a.

26. Kuramoto Y. Chemical Oscillations, Waves and Turbulence. Springer; 1983.

27. Brown E, Moehlis J, Holmes P. On the phase reduction and response dynamics of neural oscillator populations. Neural Computation. 2004;16(4):673–715. doi:10.1162/089976604322860668.

28. Rosenmund C, Stevens CF. Definition of the readily releasable pool of vesicles at hippocampal synapses. Neuron. 1996;16. doi:10.1016/S0896-6273(00)80146-4.

29. Rosenbaum R, Zimnik A, Zheng F, Turner RS, Alzheimer C, Doiron B, et al. Axonal and synaptic failure suppress the transfer of firing rate oscillations, synchrony and information during high frequency deep brain stimulation. Neurobiology of Disease. 2014;62. doi:10.1016/j.nbd.2013.09.006.

30. Hennig MH. Theoretical models of synaptic short term plasticity. Frontiers in Computational Neuroscience. 2013;doi:10.3389/fncom.2013.00045.

31. Peterson AJ, Irvine DRF, Heil P. A model of synaptic vesicle-pool depletion and replenishment can account for the interspike interval distributions and nonrenewal properties of spontaneous spike trains of auditory-nerve fibers. Journal of Neuroscience. 2014;34. doi:10.1523/JNEUROSCI.0903-14.2014.

32. Neher E. Merits and Limitations of Vesicle Pool Models in View of Heterogeneous Populations of Synaptic Vesicles. Neuron. 2015;87. doi:10.1016/j.neuron.2015.08.038.

33. Guo J, Ge JL, Hao M, Sun ZC, Wu XS, Zhu JB, et al. A Three-Pool Model Dissecting Readily Releasable Pool Replenishment at the Calyx of Held. Scientific Reports. 2015;5. doi:10.1038/srep09517.

34. Chamberland S, Tóth K. Functionally heterogeneous synaptic vesicle pools support diverse synaptic signalling. Journal of Physiology. 2016;594. doi:10.1113/JP270194.

35. Lou X, Fan F, Messa M, Raimondi A, Wu Y, Looger LL, et al. Reduced release probability prevents vesicle depletion and transmission failure at dynamin mutant synapses. Proceedings of the National Academy of Sciences of the United States of America. 2012;109. doi:10.1073/pnas.1121626109.

36. Branco T, Staras K. The probability of neurotransmitter release: Variability and feedback control at single synapses. Nature Reviews Neuroscience. 2009;10. doi:10.1038/nrn2634.

37. Qu L, Akbergenova Y, Hu Y, Schikorski T. Synapse-to-synapse variation in mean synaptic vesicle size and its relationship with synaptic morphology and function. Journal of Comparative Neurology. 2009;514. doi:10.1002/cne.22007.

38. Staras K, Branco T, Burden JJ, Pozo K, Darcy K, Marra V, et al. A Vesicle Superpool Spans Multiple Presynaptic Terminals in Hippocampal Neurons. Neuron. 2010;66. doi:10.1016/j.neuron.2010.03.020.

39. Südhof TC. The presynaptic active zone. Neuron. 2012;75. doi:10.1016/j.neuron.2012.06.012.

40. Dittman JS, Ryan TA. The control of release probability at nerve terminals. Nature Reviews Neuroscience. 2019;20. doi:10.1038/s41583-018-0111-3.

41. Duchet B, Weerasinghe G, Cagnan H, Brown P, Bick C, Bogacz R. Phase-dependence of response curves to deep brain stimulation and their relationship: from essential tremor patient data to a Wilson–Cowan model. Journal of Mathematical Neuroscience. 2020;10. doi:10.1186/s13408-020-00081-0.

42. Duchet B, Ghezzi F, Weerasinghe G, Tinkhauser G, Kühn AA, Brown P, et al. Average beta burst duration profiles provide a signature of dynamical changes between the on and off medication states in Parkinson’s disease. PLoS Computational Biology. 2021;17:e1009116. doi:10.1371/journal.pcbi.1009116.

43. Sermon JJ, Olaru M, Anso J, Little S, Bogacz R, Starr PA, et al. Sub-harmonic Entrainment of Cortical Gamma Oscillations to Deep Brain Stimulation in Parkinson’s Disease: Predictions and Validation of a Patient-Specific Nonlinear Model. bioRxiv. 2022;.

44. Schor JS, Nelson AB. Multiple stimulation parameters influence efficacy of deep brain stimulation in parkinsonian mice. Journal of Clinical Investigation. 2019;129. doi:10.1172/JCI122390.

45. Calabresi P, Ghiglieri V, Mazzocchetti P, Corbelli I, Picconi B. Levodopa-induced plasticity: A doubleedged sword in parkinson’s disease? Philosophical Transactions of the Royal Society B: Biological Sciences. 2015;370. doi:10.1098/rstb.2014.0184.

46. Steiner LA, Milosevic L. A convergent subcortical signature to explain the common efficacy of subthalamic and pallidal deep brain stimulation. Brain Communications. 2023;5. doi:10.1093/braincomms/fcad033.

47. Hashimoto T, Elder CM, Okun MS, Patrick SK, Vitek JL. Stimulation of the subthalamic nucleus changes the firing pattern of pallidal neurons. Journal of Neuroscience. 2003;23. doi:10.1523/jneurosci.23-05-01916.2003.

48. Reese R, Leblois A, Steigerwald F, Pötter-Nerger M, Herzog J, Mehdorn HM, et al. Subthalamic deep brain stimulation increases pallidal firing rate and regularity. Experimental Neurology. 2011;229. doi:10.1016/j.expneurol.2011.01.020.

49. Bergman H. The Hidden Life of the Basal Ganglia. MIT Press; 2021.

50. Realmuto J, Vidmark J, Sanger T. Modeling deep brain stimulation evoked responses with phase oscillator networks. In: 2023 11th International IEEE/EMBS Conference on Neural Engineering (NER); 2023. p. 1–4.

51. Wiest C, Torrecillos F, Tinkhauser G, Pogosyan A, Morgante F, Pereira EA, et al. Finely-tuned gamma oscillations: Spectral characteristics and links to dyskinesia. Experimental Neurology. 2022;351:113999. doi:10.1016/J.EXPNEUROL.2022.113999.

52. Cleary DR, Raslan AM, Rubin JE, Bahgat D, Viswanathan A, Heinricher MM, et al. Deep brain stimulation entrains local neuronal firing in human globus pallidus internus. Journal of Neurophysiology. 2013;109. doi:10.1152/jn.00420.2012.

53. Basu I, Crocker B, Farnes K, Robertson MM, Paulk AC, Vallejo DI, et al. A neural mass model to predict electrical stimulation evoked responses in human and non-human primate brain. Journal of Neural Engineering. 2018;15. doi:10.1088/1741-2552/aae136.

54. Detorakis G, Chaillet A, Palfi S, Senova S. Closed-loop stimulation of a delayed neural fields model of parkinsonian STN-GPe network: a theoretical and computational study. Frontiers in Neuroscience. 2015;9. doi:10.3389/fnins.2015.00237.

55. Nabi A, Moehlis J. Single input optimal control for globally coupled neuron networks. Journal of Neural Engineering. 2011;8. doi:10.1088/1741-2560/8/6/065008.

56. Gardella C, Marre O, Mora T. A tractable method for describing complex couplings between neurons and population rate. eNeuro. 2016;3. doi:10.1523/ENEURO.0160-15.2016.

57. Ozturk M, Viswanathan A, Sheth SA, Ince NF. Electroceutically induced subthalamic high-frequency oscillations and evoked compound activity may explain the mechanism of therapeutic stimulation in Parkinson’s disease. Communications Biology. 2021;4. doi:10.1038/s42003-021-01915-7.

58. Zapata Amaya V, Aman JE, Johnson LA, Wang J, Patriat R, Hill ME, et al. Low-frequency deep brain stimulation reveals resonant beta-band evoked oscillations in the pallidum of Parkinson’s Disease patient. Frontiers in Human Neuroscience. 2023;17. doi:10.3389/fnhum.2023.1178527.

59. Bolam JP, Brown MTC, Moss J, Magill PJ. Basal Ganglia: Internal Organization. Encyclopedia of Neuroscience. 2009;doi:10.1016/B978-008045046-9.01294-8.

60. Worbe Y, Yelnik J, Lehéricy S. Basal Ganglia. In: Koob GF, Moal ML, Thompson RF, editors. Encyclopedia of Behavioral Neuroscience. Oxford: Academic Press; 2010. p. 118–126. Available from: https://www.sciencedirect.com/science/article/pii/B9780080453965001706.

